# CRISPR-enabled genetic screens identify synthetic lethal targets across frequently altered cancer drivers

**DOI:** 10.64898/2026.01.25.701589

**Authors:** Jessica Desjardins, Julian Bowlan, Cynthia Bernier, Srishti Jain, Théo Goullet de Rugy, David Gallo, Mark E. Orcholski, Nancy Laterreur, Abira Rajah, Joshua Miller, Jennifer Lafontaine, Vivek Bhaskaran, Li Li, Alexanda Ling, Jesse H. Leblanc, Marie-Claude Mathieu, Michael Zinda, Stephen J. Morris, Daniel Durocher, Michal Zimmermann, Anne Roulston, Artur Veloso, Chris Fiore, Alejandro Álvarez-Quilón, Jordan T.F. Young

## Abstract

Synthetic lethality (SL) provides a treatment paradigm for targeting cancer with alterations in driver genes that are not conventionally druggable, including loss-of-function (LoF) mutations in tumor suppressor genes and gain-of-function (GoF) alterations in oncogenes. We undertook a series of genome-wide CRISPR screens using functionally validated isogenic cell lines and also conducted a large-scale SL analysis using data from the cancer dependency map (DepMap). We charted SL interactions across 15 genetic alterations characteristic of diseases with high incidence and unmet clinical need: *FBXW7*, *CCNE1*, *CDK12*, *ARID1A*, *KMT2D*, *DNMT3A*, *TET2*, *KEAP1*, *STK11*, *IDH1*, *SF3B1*, *SRSF2*, *U2AF1*, chromosome 18q loss, and chromosome 13q loss. We show validation of several SL interactions between tractable targets with cancer drivers, including *ARID1A* and the hexosamine biosynthetic pathway aminotransferase GFPT1, *STK11* with CAMK protein kinase family members including MARK2, *FBXW7* and the CDK1 regulatory kinase PKMYT1, and *CCNE1* amplification and the anaphase promoting complex or cyclosome (APC/C). In summary, this study offers a rich resource of genetic interactions across cancer drivers enabling the discovery of new biological insights and drug targets for future therapeutic development.

## Introduction

Cancer is a group of diseases characterized by uncontrolled cellular proliferation and remains one of the greatest challenges to human health. Enabled by modern multi-omics technologies, more than 350 of the approximately 22,000 protein-coding genes (about 1.6%) have been identified as cancer drivers^1^. These genes often carry alterations such as single nucleotide variants (SNVs), insertions/deletions (indels), copy number changes, rearrangements, and epigenetic silencing. While oncogenes promote cancer via GoF mutations that enhance their function, tumor suppressor genes require LoF alterations that impair their inhibitory roles for oncogenesis. Importantly, although these genetic alterations are the underlying cause of cancer, they also represent an Achilles’ heel, as they can be targeted for patient-tailored therapeutic intervention, also known as precision oncology.

There are two major conceptually distinct approaches to therapeutically leverage cancer driver genetic alterations. First, by directly inhibiting driver gene function in sustaining tumor growth (oncogene addiction), or second, by exploiting driver-specific vulnerabilities arising consequently. In the last 25 years, oncogene addiction has laid the foundation for precision oncology, with paradigmatic examples such as BCR-ABL, EGFR, and HER2-based therapies established in the clinic^2^. However, many solid tumors largely exhibit LoF mutations in tumor suppressors (e.g., *ARID1A*) or GoF alterations in traditionally undruggable oncogenes (e.g., *CCNE1*), limiting personalized therapy options. To overcome this, recent efforts in precision oncology have focused on expanding therapeutic strategies beyond inhibiting oncogene addiction by small molecules, embracing approaches such as SL^3^, antibody-based therapies^4^, targeted radiotherapies^5^, and targeted protein degradation^6^, amongst others.

SL occurs when the disruption of two genes is lethal, while inactivation of either alone is tolerated. In cancer, SL has been applied to pairs of genes where one is altered by a cancer driver mutation and the other is altered by pharmacological inhibition, eliminating cancer cells selectively^7^. In this vein, several poly(ADP-ribose) polymerase (PARP) inhibitors have been approved for the treatment of homologous recombination-deficient (HRD) cancers driven by mutations in the *BRCA1* and *BRCA2* tumor suppressor genes, clinically validating the concept of SL^8^.

In the last decade, advances in functional genomics and genome engineering have propelled the success of drug discovery pipelines supporting the identification of new SL targets and predictive biomarkers for pre-existing drugs. The ability of CRISPR-Cas9 and derived technologies such as base and prime editing to introduce precise mutations at genome scale, has enabled the identification of a second wave of high-confidence SL targets, drug candidate mechanisms of action, and repurposed drugs^9,10^. Large-scale initiatives to map genetic dependencies across hundreds of cancer cell lines were undertaken by the Broad and Sanger Institutes amongst others, through the DepMap project^11^. While effective, this approach can be limited by the scarcity of cell lines carrying less frequent driver mutations, bias towards cancer indications with more readily derivable cell lines, variable cell line-to-cell line CRISPR efficiencies, large number of co-occurring mutations, and difficulty distinguishing pathogenic from benign variants. These challenges underscore the need for complementary approaches that can precisely model specific genetic contexts. Isogenic cell line models address many of these limitations. By engineering pairs of otherwise genetically identical cell lines that differ only by a defined mutation, isogenic systems enable direct, controlled interrogation of driver-specific dependencies, including rare or understudied alterations that are underrepresented in large-scale datasets. This precision allows researchers to uncover genes essential for cell viability only in the mutant background, thereby revealing clear and causally linked SL interactions. However, target discovery in isogenic models is limited by the lack of physiological context of cancer, including the lack of co-occurring alterations and their sequence of acquisition. Sometimes, these factors could lead to the discovery of genetic interactions that do not translate from isogenic cellular models to patients^12^. Thus, before initiating therapeutic development, robust genetic target validation across many pre-clinical models with and without the cancer driver mutation is required to understand the strength and penetrance of the SL relationship. Over the past 10 years, we and others have identified, validated, and guided several next-generation SL targets including PKMYT1^13^, POLQ^14^, PLK4^15,16^, WRN^17–19^, USP1^20,21^, PRMT5, and MAT2A into the clinic^22–25^, enabling the treatment of tumors harboring traditionally un-targetable genetic alterations. Although progress has been made, many actionable SL dependencies across diverse cancer driver mutations are yet to be systematically explored.

Here, we provide a resource of genome-wide CRISPR screens across 15 isogenic cell line pairs that model cancer driver alterations associated with the cell cycle and DNA repair (*FBXW7*, *CCNE1*, and *CDK12*), chromatin regulation (*ARID1A*, *KMT2D*, *DNMT3A*, and *TET2*), cell stress and metabolism (*KEAP1*, *STK11*, and *IDH1*), alternative mRNA splicing (*SF3B1*, *SRSF2*, and *U2AF1*), and arm-level partial aneuploidy (chromosome 18q loss and chromosome 13q loss). In addition, we include analysis from DepMap tumor cell line panel screens for 14 of these cancer drivers. After comparing DepMap and isogenic screen data analysis, we demonstrate the identification of both overlapping and non-overlapping SL hits. We show the validation of one overlapping SL hit, *PKMYT1* for FBXW7-deficient cells, but also the validation of several novel isogenic screen-specific SL hits including *GFPT1* for *ARID1A*-mutated cell lines, *MARK2* for *STK11* LoF cell lines, and *UBE2C/UBE2S* for *CCNE1*-amplified cell lines.

## Results

### Genome-wide CRISPR screens reveal SL interactions across 15 cancer driver mutations

To uncover alteration-specific dependencies in cancer, we performed dropout CRISPR-based SL screens across 15 isogenic pairs representing high-prevalence driver alterations associated with significant unmet medical need (Figure S1A, B). The selected alterations encompass diverse functional classes of oncogenic events, including DNA repair and cell cycle regulation (*CCNE1, CDK12, FBXW7*), chromatin remodeling (*ARID1A, KMT2D*), cell stress response and metabolism (*KEAP1, STK11, IDH1, DNMT3A, TET2*), and mRNA alternative splicing (*SF3B1, SRSF2, U2AF1*), as well as arm-level aneuploidies frequently observed in solid tumors (18q and 13q).

Isogenic pairs were generated using CRISPR-Cas9–mediated knockout, knock-in of specific hotspot mutations, or large deletions for arm-level aneuploidies. A detailed description of all 15 isogenic pairs, including the engineering strategy, is provided in Supplementary Table 1. All models were validated through multiple orthogonal approaches, including sequencing, immunoblotting, and functional assays confirming known SL interactions, downstream target signaling, or splicing alterations, as well as whole-genome sequencing for arm-level aneuploidies (Figures S2–4).

CRISPR screens were carried out with the genome-wide TKOv3^26^ single-guide (sg)RNA library and individual sgRNA abundance was analyzed using CRISPR Count Analysis (CCA) and BAGEL2^27,28^, comparing samples obtained at the initial time point (T_0_) with their corresponding end points (T_18-21_). As expected, all screens showed strong read count correlations at T_0_, strong clustering of cell line backgrounds by principal component analysis (PCA), and a significant dropout of sgRNAs targeting common essential genes (Figure S5A-C). To identify potential SL targets, gene essentiality occurring only in genetic driver-mutant cell lines was determined by CCA and differential BAGEL2 relative to their respective parental cell lines (Table 1). Raw sgRNA read counts for all CRISPR screens reported in this study are deposited in Mendeley Data (https://doi.org/10.17632/k6wm46g4tw.1). As expected, we identified known SL interactions between *ARID1A* and its paralog *ARID1B*^29^, *CDK12* and *CDK13*^30^, *VPS4A* in cell lines that have partial arm-level aneuploidy for chromosome 18q containing its paralog *VPS4B*^31^, and *PKMYT1* in *CCNE1*-amplification^13^, as well as oncogene addiction interactions, such as the dependency on *NFE2L2* (*NRF2*) of *KEAP1*-mutated cells^32^, validating our approach (Figure 1A). Importantly, the log-transformed fold-change values of the top 20 mutation-specific dependent genes had similar magnitudes to common essential genes (Figure S5D). We observed minimal overlap in the top 50 hits across screens, with only 24 genes appearing as hits in more than two screens (Figure S5E), suggesting this approach, identified a set of unique targets for each genetic alteration.

**Figure 1.**
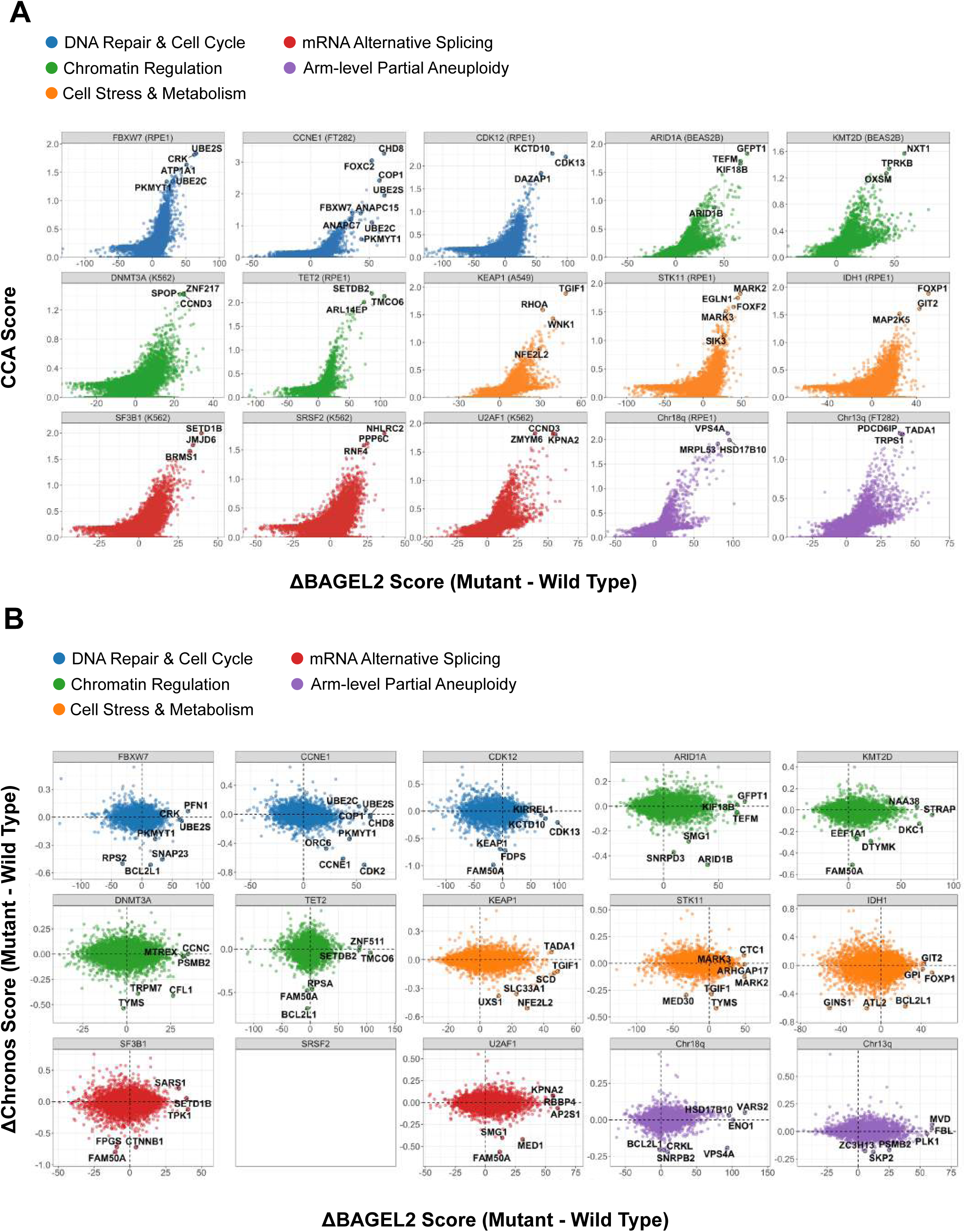
CRISPR-enabled genetic screens identify synthetic lethal interactions across frequently altered cancer drivers. **A)** Results of 15 isogenic synthetic lethal screens in the indicated cellular models carrying a single cancer driver alteration with CCA and ΔBAGEL2 (Mutant - WT) scores plotted. BAGEL2 and CCA scores both positively reflect synthetic lethal interactions. **B)** Comparison of ΔBAGEL2 scores from synthetic lethal screens in isogenic models and ΔChronos scores in cancer cell lines from the DepMap project grouped according to genetic alteration.

To compare our isogenic approach with the DepMap cancer-derived cell line dependency dataset, we identified and categorized tumor cell lines harboring pathogenic mutations in the driver genes listed above utilizing DNA sequencing data from the Cancer Cell Line Encyclopedia (CCLE)^33^. Except for *SRSF2*, we were able to identify at least three cancer cell lines that harbored pathogenic mutations in our selected driver genes. Next, we determined the differential Chronos score between wild-type and mutant cancer cell lines^34^ and compared it to the differential BAGEL2 obtained from isogenic screens for each cancer driver gene (Table 1 and Figure 1B). A list of the cancer cell lines used for this analysis can be found in Supplementary Table 2. As expected, we observed overlapping genetic interactions across the 14 cancer drivers using both methods, including *ARID1A-ARID1B*, *CDK12-CDK13*, *KEAP1-NFE2L2*, chromosome 18q-VPS4A and *CCNE1*-*PKMYT1*). In addition, we identified PKMYT1 as a SL target in both *FBXW7* isogenic screens and DepMap, supporting PKMYT1 inhibition as a potential therapeutic strategy for cancers with *FBXW7* mutations. We also identified unique SL hits using isogenic screens, including *GFPT1* for *ARID1A* LoF, *MARK2* and *MARK3* for *STK11* loss, and *UBE2C* and *UBE2S* for *CCNE1*-amplification. In summary, conducting genome-scale CRISPR screens in both isogenic and cancer cell line models represents a powerful and complementary approach for informing potential therapeutic strategies based on SL. Below, we validate and characterize a number of genetic interactions, focusing on those that are not readily detectable in DepMap.

### ARID1A deficiency creates a pronounced dependency on GFPT1 glutaminase activity

We identified the GFPT1 aminotransferase (also known as GFAT1) as one of the top SL hits in the BEAS-2B *ARID1A* isogenic pair (Figure 1A). GFPT1 is a ubiquitously expressed enzyme that catalyzes the rate-limiting step of the hexosamine biosynthetic pathway, producing 5’-diphospho-N-acetyl-glucosamine (UDP-GlcNAc) for protein glycosylation, while also contributing to energy metabolism through the conversion of glutamine to glutamate^35^. To validate the *ARID1A-GFPT1* SL interaction, we performed clonogenic survival assays upon introduction of sgRNAs targeting *GFPT1* into the BEAS-2B *ARID1A* isogenic pair (Figure 2A). *GFPT1* inactivation induced a strong selective lethality in ARID1A-deficient cells, to a similar extent to the inactivation of *ARID1B*. Metabolic cancer dependencies are often influenced by cell culture conditions and may not reliably translate to more physiologically relevant *in vivo* contexts^36^. To investigate this possibility, we performed clonogenic survival assays using a more physiologically relevant human plasma-like media (HPLM)^37^. We found that *GFPT1* sgRNAs induced lethality in *ARID1A* knockout cells to an equal degree in HPLM media compared to normal DMEM media (Figure S6A).

**Figure 2.**
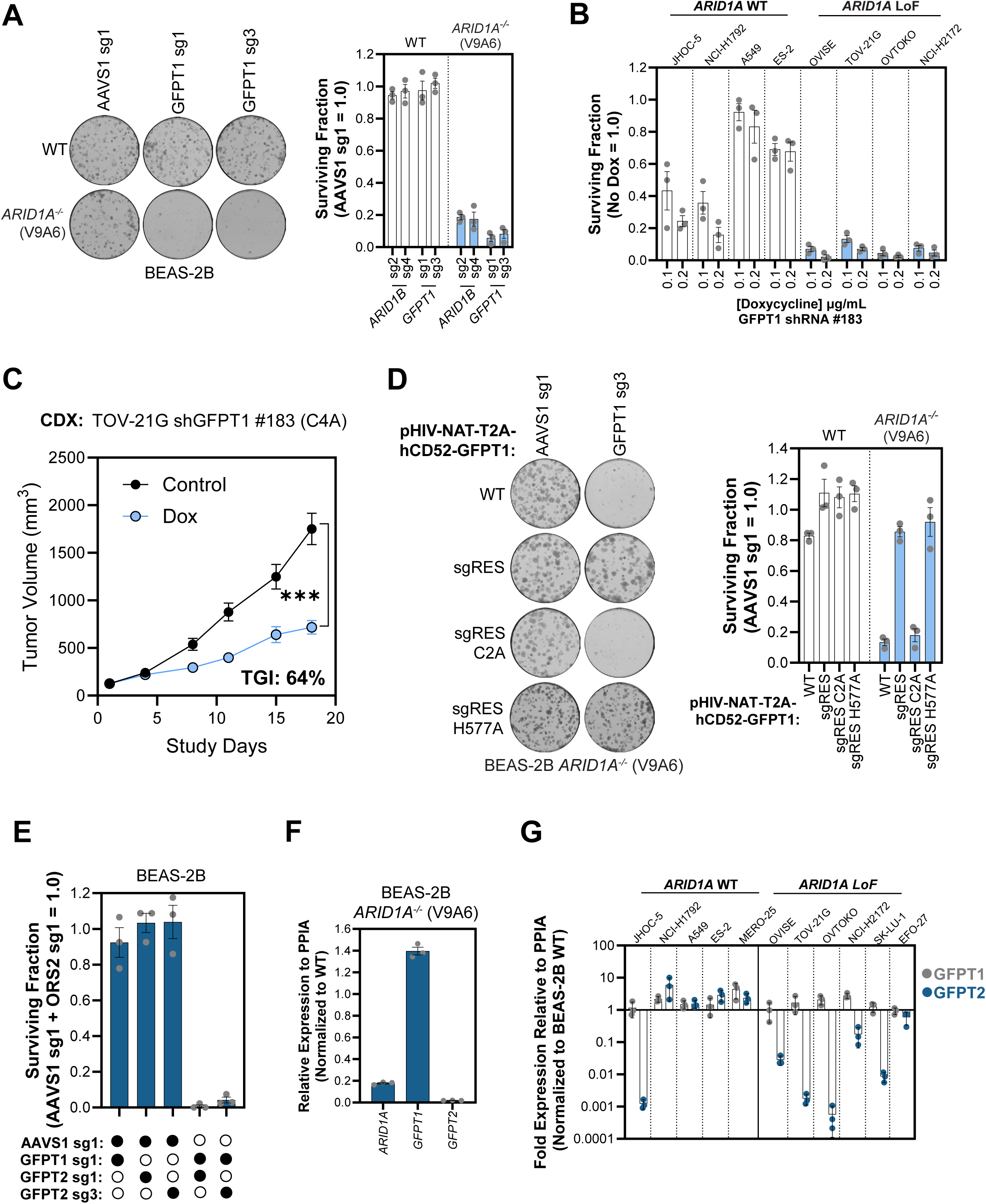
GFPT1 glutaminase activity is required for the survival of ARID1A-deficient cells. **A)** Clonogenic survival assays in BEAS-2B cells with the indicated *ARID1A* genotype upon introduction of sgRNAs targeting either *AAVS1*, *ARID1B*, or *GFPT1*. Left, representative images of colonies stained with crystal violet. Right, quantification of clonogenic survival assays. Data are the mean ± SD (*n* = 3). **B)** Clonogenic survival assays of the indicated tumor-derived cell lines upon transduction with vectors conditionally expressing a shRNA targeting *GFPT1* (shRNA#183) upon exposure to doxycycline. **C)** Tumor growth of TOV-21G cells conditionally expressing *GFPT1* shRNA#183 upon doxycycline exposure. TOV-21G cells were implanted to generate tumor xenografts in NCG mice that were randomized to receive either a vehicle control or 2 g/kg doxycycline in chow *ad libitum*. Results are expressed as mean tumor volume ± SEM (*n* = 8 mice per group). TGI and *p* value relative to vehicle was determined by unpaired one-sided *t*-test. **D)** Clonogenic survival assays in BEAS-2B *ARID1A*^-/-^ cells expressing sgRNA-resistant (sgRES) ORFs of either wild-type, glutaminase-dead (C2A), or isomerase-dead (H577A) GFPT1. The indicated mutants were nucleofected with GFPT1 sgRNA #3. Left, representative images of colonies stained with crystal violet. Right, quantitation of clonogenic survival assays. Data are represented as the mean ± SD (*n* = 3). **E)** Quantification of clonogenic survival assays in BEAS-2B cells nucleofected with combined sgRNAs targeting *AAVS1/OSR2*, *AAVS1/GFPT1*, *AAVS1*/*GFPT2,* or *GFPT1*/*GFPT2*. **F)** Relative gene expression (normalized to *PPIA*) measured by RT-qPCR for *ARID1A*, *GFPT1* and *GPFT2* transcripts in BEAS-2B *ARID1A^-/-^* cells. **G)** Relative gene expression (normalized to *PPIA*) measured by RT-qPCR for *GFPT1* and *GPFT2* transcripts in the indicated tumor-derived cell lines. Data are represented as the mean ± SD (*n* = 3).

To further test this genetic interaction in additional cellular backgrounds, we assembled a panel of tumor-derived cell lines that were either *ARID1A* wild-type (JHOC-5, NCI-H1792, A549, and ES-2) or *ARID1A* LoF (OVISE, TOV-21G, OVTOKO, and NCI-H2172). ARID1A status was confirmed by immunoblotting protein extracts from the assembled tumor cell line panel (Figure S7D). We performed clonogenic survival assays upon nucleofection of Cas9 and a *GFPT1* sgRNA (Figure S6B). Except for BEAS-2B and ES-2, *GFPT1* sgRNAs induced a strong fitness defect across our panel of tumor cell lines. Since complete *GFPT1* knockout by CRISPR was lethal in most cell lines, we hypothesized that partial depletion could result in a detectable SL window between wild-type and ARID1A-deficient cell lines. To test this possibility, we introduced two independent tetracycline-inducible *GFPT1* shRNAs by lentiviral transduction into the BEAS-2B *ARID1A* isogenic pair (Figure S6C). We observed a doxycycline-dependent knockdown of *GFPT1* expression with both shRNA #183 and #864 in wild-type and *ARID1A^-/-^* cell lines. We also detected a robust doxycycline dose-dependent clonogenic lethality in BEAS-2B *ARID1A* knockout cells compared to wild-type controls (Figure S6D).

Next, we examined whether the lethality in *ARID1A*-deficient cells was a consequence of on-target depletion of *GFPT1* by shRNA #183. To test this, we transduced the BEAS-2B isogenic pair with a lentiviral construct that expressed an shRNA-resistant *GFPT1* (Figure S6E). Consistent with an on-target effect of GFPT1 depletion by shRNA, we observed a complete rescue of the lethality. We next assessed the effect of partial inactivation of *GFPT1* with shRNA in the *ARID1A* wild-type and mutant tumor-derived cell line panel (Figure 2B). We found that *ARID1A* LoF tumor cell lines were more sensitive to *GFPT1* depletion upon doxycycline treatment compared to *ARID1A* wild-type cell lines by clonogenic survival assay. In contrast to complete knockout by CRISPR, we observed a considerable SL window with partial depletion of GFPT1 by RNAi. However, two *ARID1A* wild-type tumor cell lines, JHOC-5 and NCI-H1792, also had a sizable growth defect upon *GFPT1* partial knockdown, suggesting that additional mechanisms to ARID1A loss operate to confer dependency on GFPT1 in cancer cells.

To validate *GFPT1-ARID1A* SL *in vivo*, we established a sub-clonal cell line of the ovarian tumor cell line TOV-21G (Clone 4A) expressing doxycycline inducible shRNA#183. This cell line was implanted to generate tumor xenografts in NCG mice that were randomized to receive either a vehicle control or 2 g/kg doxycycline in chow *ad libitum*. Over the course of the study, we observed robust tumor growth inhibition (TGI = 64%; *n* = 6 mice per group) in the doxycycline fed cohort compared to the vehicle control with no effect on body weight (Figure 2C and Figure S7A). We also detected substantial *GFPT1* knockdown on study days 3, 6, and 9 by immunoblotting proteins extracted from tumors in doxycycline fed animals (Figure S7B). Together, these data suggest that *GFPT1-ARID1A* SL is not an artifact of 2D culture conditions but rather a robust genetic interaction that can be translated to *in vivo* settings.

The GFPT1 aminotransferase contains two functional domains: the isomerase domain responsible for the catalysis of fructose-6-phosphate to glucosamine-6-phosphate, and the glutaminase domain responsible for releasing ammonia in the hydrolysis of glutamine to glutamate^35,38^. The isomerase function is critical in the hexosamine biosynthetic pathway for the production of UDP-GlcNAc and subsequent protein glycosylation. In contrast, the glutaminase function is critical for providing cells with sufficient levels of glutamate required for the tricarboxylic acid (TCA) cycle and as a key building block for the synthesis of nucleotides and amino acids. We next examined which GFPT1 enzymatic activities were required for the selective lethality observed in ARID1A-deficient cells (Figure 2D). Reintroduction of an sgRNA-resistant *GFPT1* transgene rescued the lethality induced by *GFPT1* sgRNA #3 in BEAS-2B *ARID1A^-/-^*(V9A6) cells, demonstrating that the phenotype was a result of on-target CRISPR activity. Exogeneous expression of the isomerase-dead GFPT1^H577A^ mutant also protected ARID1A-deficient cells from lethality induced by endogenous *GFPT1* knockout, whereas expression of the glutaminase-dead GFPT1^C2A^ mutant did not. We conclude that the glutaminase activity of GFPT1 is critical for the survival of ARID1A-deficient cells where the isomerase activity is dispensable. These data highlight the importance of cellular glutamate production by GFPT1 rather than protein glycosylation by the hexosamine biosynthetic pathway.

Previously published CRISPR-based dual sgRNA paralog screens reported a SL interaction between *GFPT1* and its close gene paralog *GFPT2*^39^. The *GFPT1* and *GFPT2* paralogs demonstrate 80% amino acid sequence identity in humans and have similar isomerase and glutaminase activities^38^. To confirm the *GFPT1-GFPT2* genetic interaction, we introduced *GFPT1* and *GFPT2* sgRNAs into BEAS-2B wild-type cells in combination with an *AAVS1* negative control sgRNA or in combination with each other to induce single or double knockouts, respectively (Figure 2E). Clonogenic survival assays showed strong lethality in cells nucleofected with both *GFPT1* and *GFPT2* sgRNAs compared to single knockouts, confirming the reported paralog SL. Since ARID1A is a component of SWI/SNF remodeling complexes and can activate or repress transcription of select genes, we hypothesized that *ARID1A*-deficiency could impact *GFPT2* gene expression resulting in a dependency on *GFPT1*. To test this possibility, we extracted RNA from BEAS-2B wild-type and *ARID1A^-/-^* (V9A6) cells and performed quantitative reverse transcription polymerase chain reaction (RT-qPCR) using specific probes for *ARID1A*, *GFPT1*, and *GFPT2* (Figure 2F). We observed a dramatic loss of *GFPT2* mRNA expression in *ARID1A* knockout cells compared to wild-type controls. Furthermore, very low GFPT2 protein levels were also detected in BEAS-2B ARID1A-deficient cells compared to wild-type (Figure S7C). In contrast, there was no detectable impact of ARID1A loss on *GFPT1* mRNA and protein levels. To examine *GFPT2* mRNA expression in different ARID1A-deficient cell backgrounds, we conducted RT-qPCR across our assembled tumor-derived cell line panel (Figure 2G). We included one additional *ARID1A* wild-type tumor cell line (MERO-25) and two additional *ARID1A* LoF cell lines (SK-LU-1 and EFO-27). We found that all six ARID1A-deficient tumor cell lines had lower levels of *GFPT2* mRNA compared to wild-type cells. Only one *ARID1A* wild-type tumor cell line, JHOC-5, displayed lower *GFPT2* mRNA levels. These *GFPT2* mRNA expression results were also confirmed at the protein level by immunoblotting (Figure S7D). As expected, JHOC-5 cells were also sensitive to *GFPT1* shRNA-mediated knockdown (Figure 2B). Importantly, one *ARID1A* wild-type tumor cell line, NCI-H1792, had normal *GFPT2* mRNA expression but was sensitive to *GFPT1* knockdown, suggesting that *GFPT2* expression may not be the only determinant of *GFPT1* dependency in tumor cell lines. Together, these data suggest that *ARID1A*-deficiency is associated with lower *GFPT2* gene expression, resulting in a requirement for *GFPT1* for cellular survival. Since the two paralogs are SL, future therapeutic development should focus on selectively targeting GFPT1 with minimal off-target activity on GFPT2 to avoid unwanted toxicity in normal tissues.

### The tumor suppressor gene *STK11* is synthetic lethal with the MARK2 kinase

We performed genome-wide CRISPR screens in the RPE1-hTERT Cas9 *p53^-/-^ STK11* isogenic pair and identified several CAMK kinase genes as top hits including *MARK2* and *MARK3*, members of the microtubule affinity-regulating kinase (MARK) family, and *SIK3,* a member of the salt-inducible kinase (SIK) family (Table 1 and Figure 1A). In addition to the role of STK11 in phosphorylating the AMPKα activation loop in conditions of energy stress, *STK11* has been shown to phosphorylate MARK and SIK kinases^40^, suggesting a functional link between them. To validate the *STK11-MARK* SL interaction, we carried out clonogenic survival assays introducing two independent *MARK2* or *MARK3* targeting sgRNAs into wild-type RPE1-hTERT Cas9 *p53^-/-^*or *STK11* knockout cells (Figure 3A). We observed that inactivation of *MARK2* or *MARK3* induced a selective growth defect in STK11-deficient cells compared to wild-type controls. Since MARK2 and MARK3 are known to phosphorylate similar substrates, we hypothesized that combined loss could enhance the SL window between STK11-deficient cells and their parental counterparts. To test this possibility, we introduced both *MARK2* and *MARK3* targeting sgRNAs in combination. Contrary to our prediction, we observed dramatic cell death in both wild-type and *STK11* knockout cells when *MARK2* and *MARK3* were simultaneously inactivated. In support of these findings, we found that *MARK2-MARK3* paralog dependency was previously observed in multiple tumor cell line backgrounds using combinatorial sgRNA CRISPR screens^41^. We next examined whether the kinase activity of MARK2 was required for the SL observed in STK11-deficient cells. For this, we reintroduced sgRNA-resistant wild-type MARK2 or a kinase-dead MARK2^K82M^ mutant^42^ in RPE1-hTERT Cas9 *p53^-/-^ STK11^-/-^*(C19) cells (Figure 3B). While expression of wild-type MARK2 protected STK11-deficient cells from cell death, the kinase-dead MARK2^K82M^ mutant did not. We conclude that the kinase activity of MARK2 is critical for promoting survival of STK11-deficient cells.

**Figure 3.**
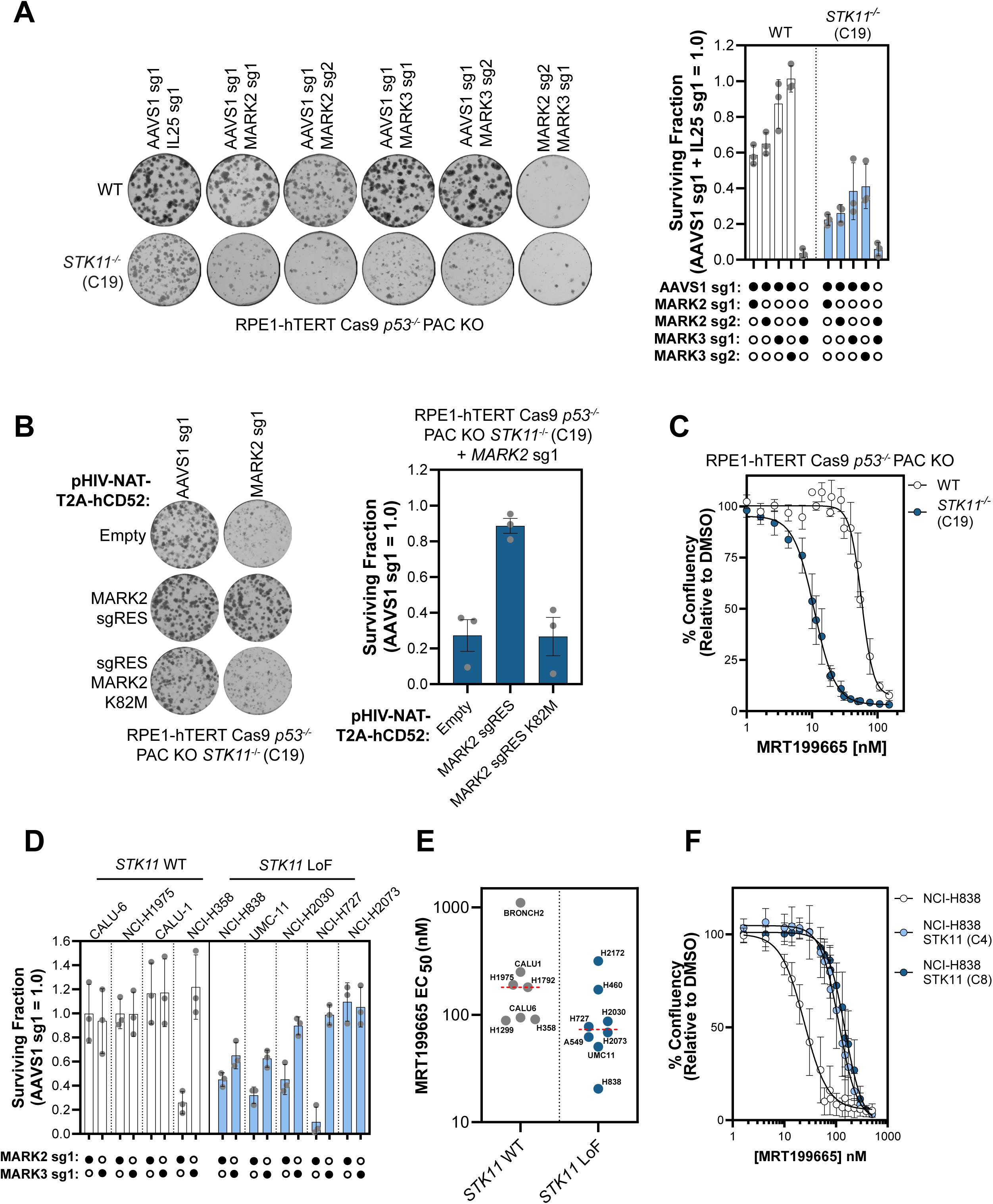
MARK2 kinase is synthetic lethal with STK11. **A)** Clonogenic survival assays in RPE1-hTERT Cas9 *p53^−/−^* cells with the indicated *STK11* genotype after introduction of combined sgRNAs targeting *AAVS1/IL25*, *AAVS1/MARK2*, *AAVS1*/*MARK3*, or *MARK2*/*MARK3*. Left, representative images of colonies stained with crystal violet. Right, quantification of clonogenic survival assays. Data are the mean ± SD (*n* = 3). **B)** Clonogenic survival assays of RPE1-hTERT Cas9 *p53^−/−^ STK11^-/-^* cells expressing sgRNA-resistant (sgRES) wild-type or kinase-dead (K82M) MARK2 after introduction of sgRNA #1 targeting endogenous *MARK2*. Left, representative images of colonies stained with crystal violet. Right, quantitation of clonogenic survival assays. Data are represented as the mean ± SD (*n* = 3). **C)** Dose-response proliferation assays with MRT199665 in RPE1-hTERT Cas9 *p53^−/−^* cells with the indicated *STK11* genotype. Data are represented as mean ± SD (*n* = 3). **D)** Clonogenic survival assays in the indicated tumor-derived cell lines after the introduction of Cas9 and an sgRNA targeting *AAVS1*, *MARK2*, or *MARK3*. The mutation status of the *STK11* gene is indicated. Data are the mean ± SD (*n* = 3). **E)** Dose-response proliferation assays with MRT199665 in the indicated tumor-derived cell lines. EC₅₀ values are presented as the mean of three biological replicates. Median values for wild-type and *STK11*-mutant groups are shown as red dotted lines. **F)** Dose-response proliferation assays with MRT199665 in NCI-H838 lung cancer-derived parental cells and two independent *STK11* transgene overexpressing clones (C4 and C8). Data are represented as the mean ± SD (*n* = 3).

Since multiple CAMK kinase family members were hits in the *STK11* SL screen, including *MARK2*, *MARK3*, and *SIK3*, we hypothesized that pharmacological inhibition of these kinases could result in a selective dependency in STK11-deficient cells. To investigate this further, we obtained a previously reported small molecule tool called MRT199665 which is a potent but non-selective kinase inhibitor of several CAMK family members including AMPKα, MARK, SIK, NUAK, and MELK^43^. Importantly, MRT199665 does not selectively target the STK11 tumor suppressor kinase, making it a suitable pharmacological tool in this context. We performed dose-response proliferation assays in the *STK11* isogenic pair treated with MRT199665 (Figure 3C). We observed a 5.9-fold increase in sensitivity to MRT199665 compared to wild-type, confirming the dependency of STK11-deficient cells on CAMK kinases. However, wild-type cells also displayed significant growth inhibition in response to MRT199665 (EC_50_ value of 63.6 nM), suggesting that combined inhibition of multiple CAMK kinases may be generally lethal to cells.

To test the *STK11-MARK* genetic interaction in additional backgrounds, we assembled a panel of tumor-derived cell lines that were either *STK11* wild-type (CALU-6, NCI-H1975, CALU-1, and NCI-H358) or *STK11* LoF (NCI-H838, UMC-11, NCI-H2030, NCI-H727, and NCI-H2073). STK11 status was confirmed by immunoblotting protein extracts from the assembled tumor cell line panel (Figure S7E). For each tumor cell line, we introduced both Cas9 and either a *MARK2* or *MARK3* sgRNA followed by seeding in clonogenic survival assays (Figure 3D). We observed *MARK2* dependency in most STK11-deficient tumor cell lines, except for NCI-H2073. Importantly, the *MARK2* sgRNA did not induce lethality in most *STK11* wild-type tumor cell lines, except for NCI-H358. In addition, we observed a significant, but milder and less penetrant (2 out of 5) dependency for *MARK3* in STK11-deficient tumor cell lines. Next, we tested the non-selective CAMK kinase inhibitor MRT199665 across our assembled tumor cell line panel using IncuCyte proliferation assays (Figure 3E). We determined that STK11-deficient tumor cells were more sensitive to MRT199665 compared to wild-type cells. However, the difference between the average EC_50_ values for both groups was small, and the effect did not translate to all tumor cell lines. To determine the contribution of STK11 loss to the sensitivity induced by MRT199665, we reintroduced a *STK11* transgene into the most sensitive tumor cell line, NCI-H838 (Figure 3F). After lentiviral transduction, we isolated two single cell clones, C4 and C8, that overexpress *STK11* as demonstrated by immunoblotting (Figure S7F). We found that exogenous expression of *STK11* in NCI-H838 cells resulted in only a 5-fold resistance to MRT199665 treatment and not complete rescue. Therefore, STK11 status is not the only determinant that can predict sensitivity to MRT199665. Together, our data suggest that *STK11* LoF tumor cell lines are more dependent on the MARK kinases compared to wild-type cell lines, however the phenotype is not fully penetrant across all cell line backgrounds and is stronger for MARK2 compared to MARK3.

### The *FBXW7* tumor suppressor is synthetic lethal with the CDK1 inhibitory kinase PKMYT1

Both our isogenic approach and DepMap cancer cell line dependencies converged in identifying the PKMYT1 kinase (also known as Myt1) as one of the top SL targets in cells with *FBXW7* mutations (Table 1 and Figure 1B). Previously, we reported a robust SL interaction between *CCNE1* amplification and *PKMYT1*^3^ and described the development of RP-6306 (lunresertib), a highly potent and selective PKMYT1 small molecule inhibitor^44^. To examine the *FBXW7-PKMYT1* SL relationship further, we performed a genome-wide chemical genetic screen using RP-6175, a lunresertib analog (Figure 4A and Supplementary Table 3)^44^. Consistent with our SL CRISPR screen in *FBXW7* isogenic cells, we identified *FBXW7* as a hit, demonstrating further that disruption of *FBXW7* sensitizes cells to PKMYT1 inactivation. We also identified regulators of protein phosphatase 2A (PP2A) including *LCMT1*, *PTPA*, and *TIPRL*, hinting that oncogenic mutations impacting PP2A activity observed in aggressive uterine cancers could be sensitive to PKMYT1 inhibition^45^. Finally, to confirm the *FBXW7-PKMYT1* SL interaction, we performed clonogenic assays in the RPE1-hTERT Cas9 *p53^-/-^ FBXW7^-/-^* isogenic model. Introduction of sgRNAs targeting *PKMYT1* was selectively lethal in *FBXW7* knockout cells but not in wild-type controls (Figure 4B). All gene editing efficiencies in this study were examined by Sanger Sequencing and ICE analysis software (https://ice.editco.bio/) and can be found in Supplementary Table 7. In cancer patients, *FBXW7* mutations often manifest as biallelic truncating mutations or as monoallelic missense hotspot mutations in arginine 465 (R465), 479 (R479) or 505 (R505)^46^. These three amino acids are in the WD40 repeats and are critical for substrate recognition and CCNE1 degradation. To probe at scale which *FBXW7* mutations result in sensitivity to PKMYT1 inhibition, we undertook a base editing screen using DLD-1 cells expressing the FNLS cytosine base editor (CBE)^47^ and a focused library of sgRNAs spanning the *FBXW7* coding sequence (Figure 4C, Figure S8A, and Supplementary Table 4). Several missense variants including the well-known oncogenic driver R505C sensitized cells to PKMYT1 inhibition. In addition, we identified several novel codons that are less commonly mutated in tumors including His420, Arg441, Ser476, Asp487, and Gly644 that also conferred sensitivity to PKMYT1 inhibition. Finally, we observed nonsense truncating mutations that conferred PKMYT1i sensitivity, as expected.

**Figure 4.**
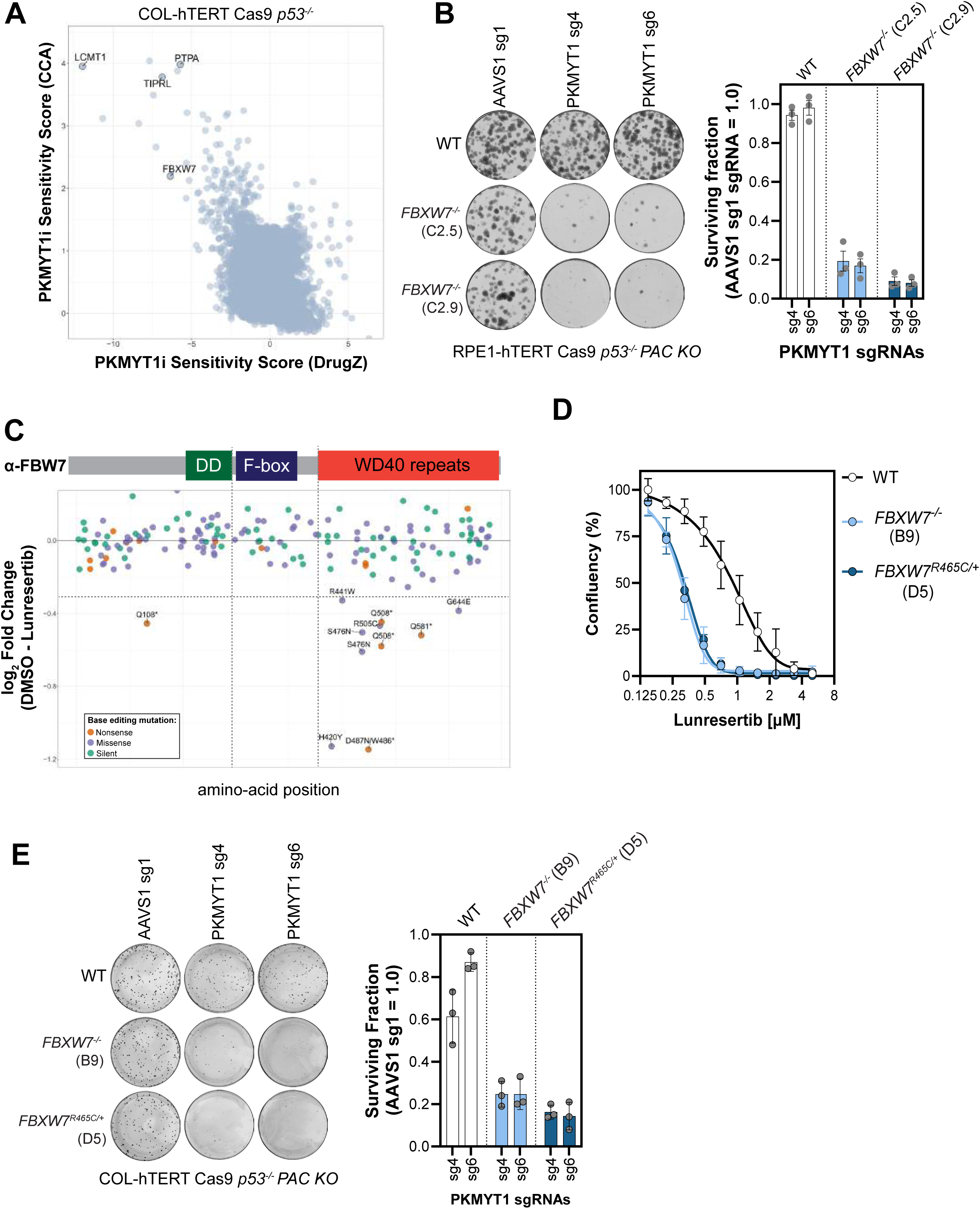
Identification of *PKMYT1-FBXW7* synthetic lethality. **A)** PKMYT1 inhibitor (RP-6175) dropout CRISPR screen performed in COL-hTERT Cas9 *p53^-/-^* cells with DrugZ and CCA scores plotted **B)** Clonogenic survival assays in RPE1-hTERT Cas9 *p53^−/−^* cell lines with the indicated *FBXW7* genotypes after introduction of sgRNAs targeting *AAVS1* or *PKMYT1*. Left, representative images of colonies stained with crystal violet. Right, quantification of clonogenic survival assays. Data are the mean ± SD (*n* = 3). **C)** Lunresertib cytosine base editing dropout screen performed in DLD-1 FNLS clone 9 cells using a focused sgRNA library targeting the *FBXW7* gene. Log2 fold-change of sgRNA effects in PKMYT1i-treated versus vehicle (DMSO) conditions, along with the predicted mutation induced by each sgRNA are shown. **D)** Dose-response proliferation assays with lunresertib in COL-hTERT Cas9 *p53^−/−^* cells with the indicated *FBXW7* genotypes. Data are represented as the mean ± SD (*n* = 3). **E)** Clonogenic survival assays in COL-hTERT Cas9 *p53^−/−^* cell lines with the indicated *FBXW7* genotypes after introduction of *AAVS1* or *PKMYT1* sgRNAs. Left, representative images of colonies stained with crystal violet. Right, quantification of clonogenic survival assays. Data are the mean ± SD (*n* = 3).

To compare the effect of PKMYT1 inhibition upon either truncating or hotspot mutations in *FBXW7*, we engineered *FBXW7^-/-^* and *FBXW7^R465C/+^* clonal cell lines in COL-hTERT Cas9 *p53^-/-^*cells using CRISPR-mediated knockout and knock-in, respectively (Supplementary Table 1 and Figure S2B). Two independent *FBXW7* biallelic knockouts and one monoallelic R465C clone were identified and as expected, all three clones had elevated CCNE1 and c-Myc protein levels, both known FBW7 substrates^48,49^. We performed dose-response growth curves upon PKMYT1 inhibition (Figure 4D) and clonogenic assays using sgRNAs targeting PKMYT1 (Figure 4E). Both COL-hTERT Cas9 *p53^-/-^ FBXW7* knockout and the R465C hotspot mutant were sensitive to PKMYT1 inhibition and genetic inactivation.

Next, to further validate the *FBXW7-PKMYT1* SL interaction, we assembled a tumor-derived cell line panel consisting of wild-type cell lines (NUGC-3, KYSE-30, and SW-480) and *FBXW7* mutated cell lines (EMTOKA, ESS-1, and KLE). For each cell line, we determined the EC_50_ for lunresertib using IncuCyte proliferation assays (Figure 5A). We found that cancer-derived cell lines harboring *FBWX7* mutations showed decreased cell growth upon PKMYT1 inhibition compared to *FBXW7* wild-type cell lines. To test this interaction in a more physiological context, we examined lunresertib in two patient-derived xenografts (PDX), one of head and neck squamous cell carcinoma origin and the other of stomach adenocarcinoma, with inactivating *FBXW7* mutation R479Q and a deep deletion, respectively. Twice-daily dosing of lunresertib at 20 mg kg−1 led to a dose-dependent tumor growth inhibition, reaching 73-90% at Day 28 of treatment (Figure 5B) with less than 10% body weight loss (Figure S8B). Together, these data demonstrate that inactivation of *PKMYT1* and mutations in the tumor suppressor *FBXW7* are SL across multiple cellular models, as well as *in vivo* PDX models. Importantly, this genetic interaction is penetrant across the full spectrum of clinically relevant *FBXW7* mutations, including both truncating and missense hotspot variants. These findings broaden the application of PKMYT1 inhibitors beyond the treatment of *CCNE1*-amplified tumors, potentially increasing the number of patients who may benefit from this mechanism of action.

**Figure 5.**
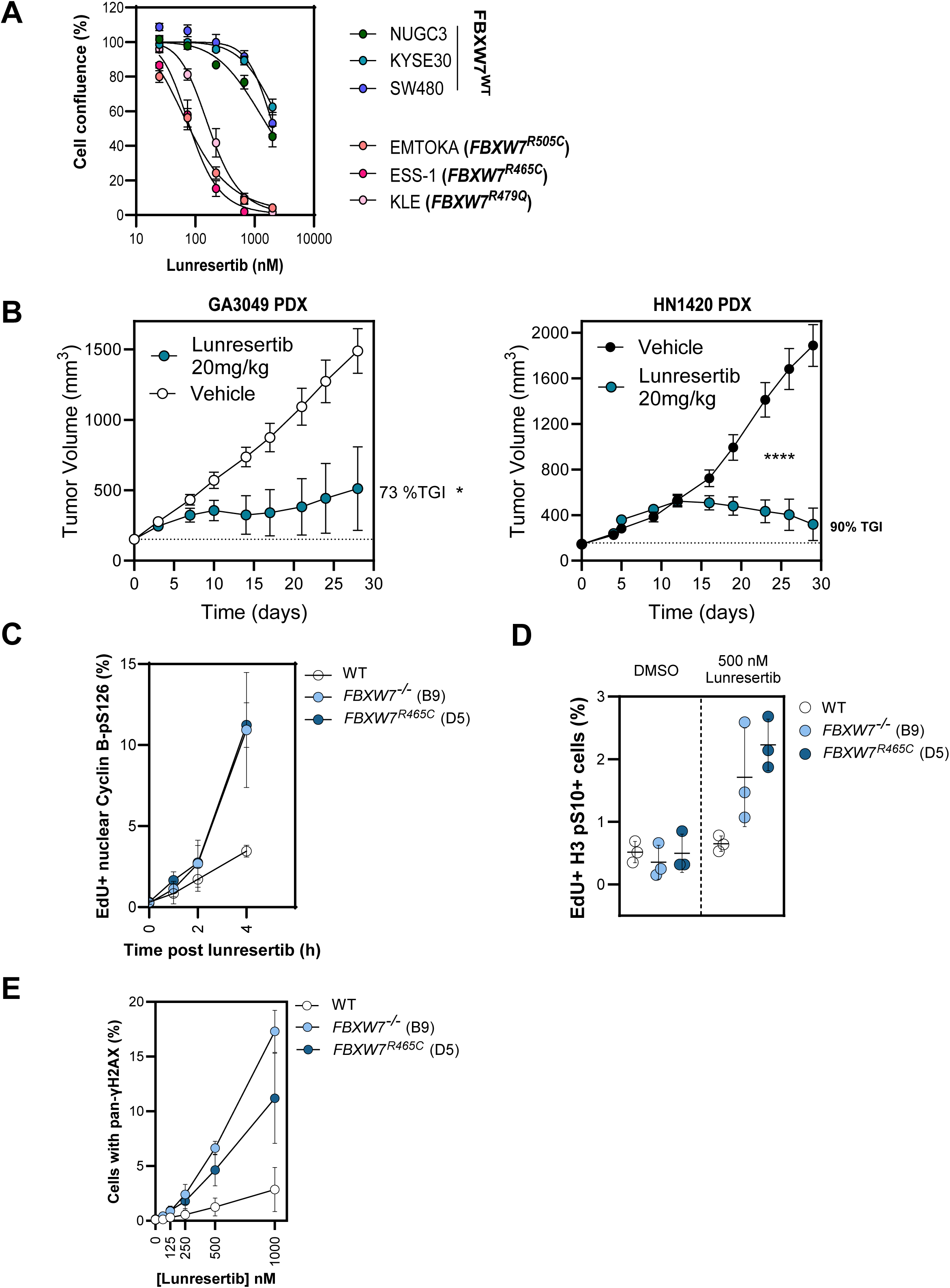
Characterization of *PKMYT1-FBXW7* synthetic lethality. **A)** Dose-response proliferation assays with lunresertib in tumor-derived cell lines with the indicated *FBXW7* genotypes. Data are represented as the mean ± SD (*n* = 3). **B)** Tumor growth of two *FBXW7*-mutated PDX models implanted in BALB/c nude mice and treated with lunresertib (20 mg kg^−1^) or vehicle administered orally twice daily for the duration of the experiment. Left, gastric cancer PDX (GA3049). Right, head and neck cancer PDX (HN1420). Results are expressed as mean tumor volume ± SEM (*n* = 8 mice/group). Statistical differences calculated using an unpaired t-test with Welch’s correction; *p<0.01, ****p<0.0001. **(C)** QIBC quantification of COL-hTERT Cas9 *p53^-/-^* cells of the indicated *FBXW7* genotype positive for EdU and cyclin B (CCNB1) pS126 positive (*n* = 3) **(D)** Flow cytometry-based quantification of COL-hTERT Cas9 *p53^-/-^* cells of the indicated *FBXW7* genotype positive for EdU and H3 pS10 (*n* = 3) with and without lunresertib treatment. **(E)** QIBC quantification of COL-hTERT Cas9 *p53^-/-^* cells of the indicated *FBXW7* genotype positive for high γH2AX, as a function of lunresertib dose (*n* = 3).

In previous work, we described the molecular basis of the SL interaction between *PKMYT1* and *CCNE1*, showing that inhibition of PKMYT1 leads to unscheduled activation of CDK1, which drives premature mitotic entry and extensive DNA damage specifically in the context of elevated CCNE1 levels^13^. Based on this, we hypothesized that LoF mutations in *FBXW7*, which are known to impair degradation of its substrate CCNE1, could result in a similar phenotype. To test whether PKMYT1 inhibition caused premature CDK1 activation in the context of FBW7 loss, we treated *FBXW7*-mutated and parental wild-type controls with lunresertib and used quantitative imaging-based cytometry (QIBC) to measure the percentage of EdU-positive cells with nuclear cyclin B1 phosphorylated at Serine 126, a marker for activated cyclin B-CDK1 complexes^41^ (Figure 5C). Upon PKMYT1 inhibition, we observed an increase in the percentage of *FBXW7*-mutated cells with EdU-positive nuclear phospho-cyclin B1 (S126) compared to wild-type, suggesting that *FBXW7* loss results in premature CDK1 activation that must be kept in check by PKMYT1 inhibitory phosphorylation. To characterize the effect of PKMYT1 inhibition on mitotic entry in *FBXW7*-mutated cells, we used flow cytometry to measure the proportion of EdU-positive cells with histone H3 phosphorylated at Ser10, a proxy for cells simultaneously undergoing DNA replication and cell division (Figure 5D). Lunresertib treatment in *FBXW7*-mutated cells induced a greater proportion of EdU-positive cells in the mitotic phase of the cell cycle compared to wild-type controls, suggesting that upon PKMYT1 inhibition FBW7 loss drives premature mitotic entry in cells still carrying out DNA replication. Finally, to test whether premature CDK1 activation and mitotic entry in *FBXW7*-mutated cells treated with lunresertib caused DNA damage, we treated the isogenic pairs with lunresertib and used QIBC to measure the percentage of cells with high pan-γH2AX intensity (Figure 5E). PKMYT1 inhibition resulted in a higher proportion of *FBXW7*-mutated cells with pan-γH2AX compared to wild-type controls, suggesting premature mitotic entry and subsequent induction of DNA damage. Together, these findings suggest that PKMYT1 safeguards unscheduled mitosis in *FBXW7*-mutated cells akin to *CCNE1*-amplification described previously.

### Oncogenic *CCNE1* amplification is synthetic lethal with UBE2C/UBE2S ubiquitin-conjugating enzymes associated with the anaphase-promoting complex

Next, we performed a genome-wide CRISPR screen using TKOv3 in both the parental FT282-hTERT *p53^R175H^* and *CCNE1*-overexpressing clone C3 cell lines and identified the PKMYT1 kinase as expected. We also found subunits of the anaphase-promoting complex/cyclosome (APC/C) including ANAPC7, ANAPC15, UBE2C, and UBE2S as top hits by CCA and BAGEL2 (Table 1 and Figure 1A). The APC/C is a multi-subunit E3 ubiquitin ligase that regulates mitotic progression by promoting the proteasomal degradation of key cell cycle proteins, working in coordination with UBE2C and UBE2S to initiate and extend Lys11-linked ubiquitin chains^50–53^. Given the essential role of these E2 enzymes in mitotic exit and existing precedence for targeting E2s with small molecules, we prioritized investigating the *CCNE1–UBE2C/S* genetic interaction. We observed that both *UBE2C* and *UBE2S* sgRNAs selectively induced lethality in *CCNE1*-overexpressing cells compared to wild-type controls in clonogenic survival assays (Figure 6A).

**Figure 6.**
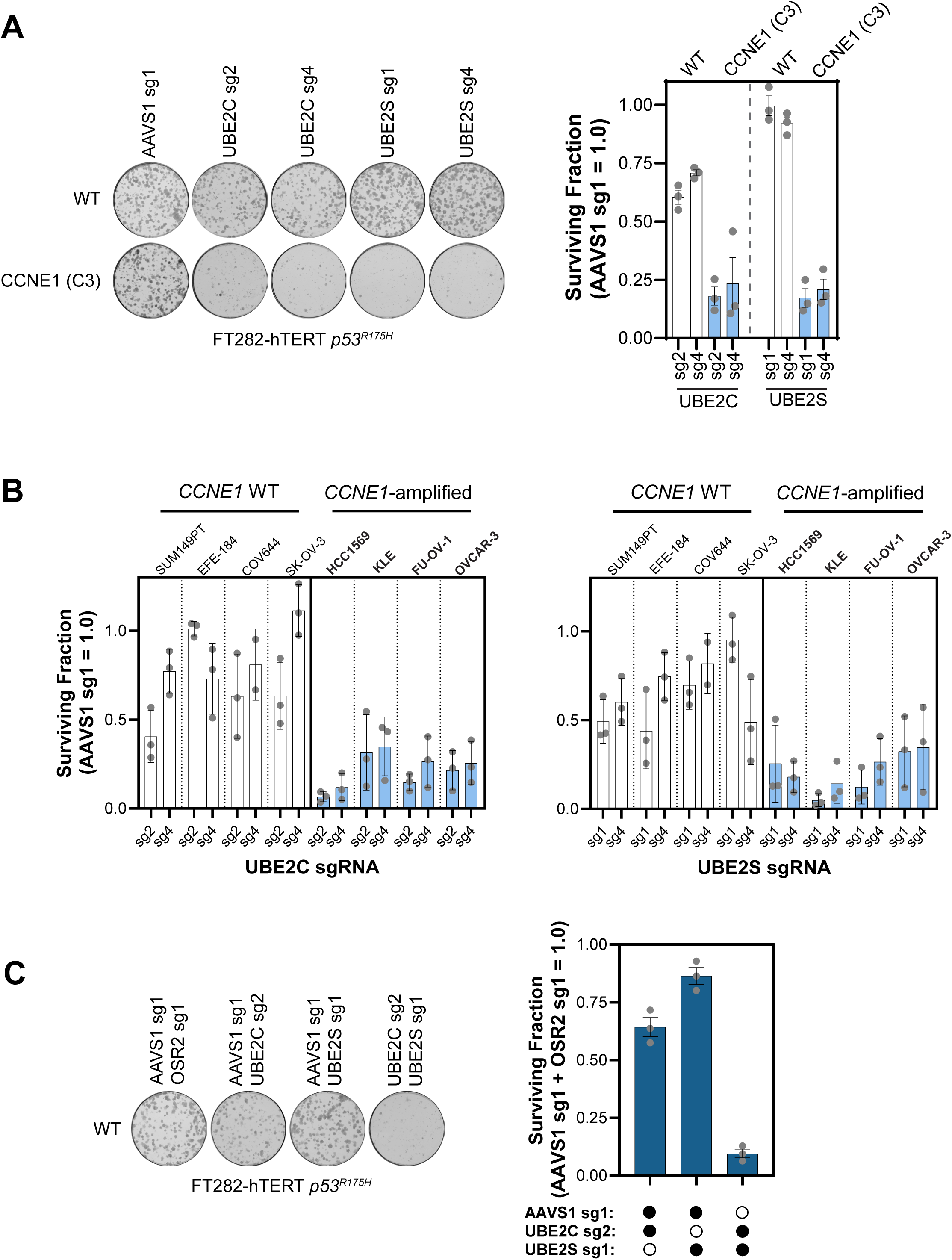
*UBE2C* and *UBE2S* are synthetic lethal with *CCNE1*-overexpression. **A)** Clonogenic survival assays in FT282-hTERT *p53^R175H^* cells upon introduction of sgRNAs targeting *AAVS1*, *UBE2C*, or *UBE2S*. Left, representative images of colonies stained with crystal violet. Right, quantification of clonogenic survival assays. Data are the mean ± SD (*n* = 3). **B)** Clonogenic survival assays of the indicated tumor-derived cell lines upon introduction of Cas9 and the indicated sgRNAs targeting *AAVS1*, *UBE2C* (left) or *UBE2S* (right). Data are represented as the mean ± SD (*n* = 3). **C)** Clonogenic survival assays in FT282-hTERT *p53^R175H^* cells after introduction of sgRNAs targeting either *AAVS1, OSR2*, *AAVS1/UBE2C* (combined), *AAVS1*/*UBE2S* (combined), or *UBE2C*/*UBE2S* (combined). Left, representative images of colonies stained with crystal violet. Right, quantification of clonogenic survival assays. Data are represented as the mean ± SD (*n* = 3).

Next, to extend validation to additional cell line backgrounds, we assembled a tumor-derived cell line panel that was either *CCNE1* wild-type (SUM149PT, EFE-184, COV644, and SK-OV-3) or *CCNE1*-amplified (HCC1569, KLE, FU-OV-1, and OVCAR-3). For each cell line, we performed clonogenic assays after introduction of Cas9-sgRNA ribonucleoprotein complexes targeting either *UBE2C* or *UBE2S* (Figure 6B). Overall, *UBE2C* and *UBE2S* sgRNAs induced a greater defect on cell fitness in *CCNE1*-amplified tumor cell lines compared to wild-type. Finally, we asked whether combined loss of both UBE2C and UBE2S would enhance the fitness defect observed in *CCNE1*-overexpressing cells. Instead, we found that combined knockout of *UBE2C* and *UBE2S* lead to complete cell death irrespective of *CCNE1* status (Figure 6C). Together, these data put forward a penetrant SL genetic interaction between *UBE2C* or *UBE2S* with *CCNE1*-amplification. Importantly, future therapeutic development should focus on selectively targeting either UBE2C or UBE2S with a small molecule inhibitor, as we discovered that the two E2 enzymes display a strong SL relationship.

## Discussion

The systematic study of cancer genetic dependencies holds promise for the development of novel targeted therapies. This work supplies a rich resource of novel SL interactions across common cancer alterations in high-prevalence cancers with unmet clinical need. Here, we provided genome-scale CRISPR screen data for 15 isogenic cell line pairs capturing major oncogenic drivers. We successfully validated several SL interactions that we discovered from these datasets including *ARID1A-GFPT1*, *STK11-MARK2*, *FBXW7-PKMYT1*, and *CCNE1-UBE2C/S*. We also compare the two major CRISPR-based approaches to SL target discovery: isogenic mutant models and large-scale cancer cell line datasets (DepMap). Although SL screens in cancer-derived cell lines have been critical to identify SL targets such as WRN, PRMT5, MAT2A, and PELO-HBS1L^17,23–25,54,55^, several hits from our isogenic approach were notably absent from DepMap analysis. These included SL interactions such as *GFPT1* (for *ARID1A* LoF)*, MARK2* (for *STK11* LoF), and *UBE2C* and *UBE2S* (for *CCNE1*-amplification), amongst others. These false negatives could reflect methodological limitations, including inaccurate LoF/GoF calls in the CCLE genomic data, insufficient number of tumor cell lines with the alteration, or heterogeneous CRISPR-Cas9 activity in tumor cell lines. These factors could contribute to a low signal-to-noise ratio in the DepMap analysis and higher false negative rate. Moreover, although 14 of 15 driver mutations modeled in isogenic backgrounds were cross-validated with DepMap, no cancer-derived models carrying oncogenic *SRSF2* mutations were identified. *SRSF2* mutations occur in ∼30–50% of chronic myelomonocytic leukemia (CMML) cases, highlighting the need to uncover novel dependencies that may inform new therapeutic strategies^56^. We propose that parallel use of cancer-derived cell lines and isogenic models provide complementary opportunities to identify novel SL drug targets. Ultimately, orthogonal validation in both isogenic and cancer-derived cell models, as well as *in vivo* studies, will be the critical gatekeeper for target selection.

We report a novel SL interaction between *ARID1A* and *GFPT1*. Importantly, we found that the glutaminase activity of GFPT1, which converts glutamine to glutamate, is essential for ARID1A-deficient cells, whereas its isomerase activity is not. We hypothesize that ARID1A-deficient cells may rely on glutamate, which is required for energy metabolism in the TCA cycle and the biosynthesis of nucleotides, amino acids, and other organic compounds such as glutathione. Notably, the importance of glutamate-derived metabolites for the survival of ARID1A-deficient tumors has been previously demonstrated. For example, ARID1A-deficient cells were shown to be vulnerable to inhibition of glutathione and the glutamate-cysteine ligase synthetase catalytic subunit, the rate-limiting enzyme for glutathione synthesis^57^. Second, another study demonstrated a dependence on cellular glutamate in ARID1A-deficient cells using a glutaminase inhibitor, CB-839, in cells, and *in vivo*^58^. Alternatively, our work also pointed to the deregulation of the *GFPT1* gene paralog, *GFPT2*, as a potential determinant of the SL in ARID1A-deficient cells. We and others have documented a strong SL relationship between the *GFPT1* and *GFPT2* paralogs^39^, regardless of the genetic background. Thus, we hypothesize that a direct downregulation of *GFPT2* expression caused by ARID1A loss may lead cells to rely on GFPT1 activity for survival. Future studies should focus on directly measuring the presence of SWI-SNF complexes at the promoter of *GFPT2.* This will help determine if the negative effect on *GFPT2* expression in ARID1A-deficient cells is a direct consequence of defective SWI-SNF transcriptional regulation. Nonetheless, our work provides cellular and *in vivo* validation of GFPT1 glutaminase activity as a tractable avenue for novel therapeutic interventions in tumors carrying mutations in the *ARID1A* tumor suppressor.

The STK11 tumor suppressor is the master kinase that activates the AMPK subfamily including the four MARK kinases^40^. In particular, the STK11-MARK2 signaling cascade has been shown to regulate many essential cellular processes including the establishment of cell polarity, cell cycle regulation, vesicular transport, and cell migration^59^. Studies have shown that STK11 can activate the kinase activity of MARK2 by approximately 200-fold through the phosphorylation of Thr208^40^. However, our data suggests that the kinase activity of MARK2 has essential cellular functions in the absence of STK11. One possibility is that another factor could activate MARK2 kinase activity in the absence of STK11. Importantly, a STE family kinase, MARKK/TAO1, was previously demonstrated to also activate MARK2 through phosphorylation of Thr208^52^. Future studies should focus on the mechanisms that promote MARK2 activation in the absence of STK11. Here, we have demonstrated for the first time that chemical or genetic inactivation of MARK2 kinase activity, and to some extent MARK3, leads to selective growth inhibition in STK11-deficient cells. One explanation for this SL interaction is that the combined effect of STK11 and MARK2 loss on optimal STK11 pathway activation falls below the required threshold for cell survival. Future work should evaluate which cellular processes determine the observed cell death in STK11-deficient cells after loss of MARK2 kinase activity.

Lunresertib (RP-6306) is a first-in-class PKMYT1 inhibitor currently in Phase 1/2 clinical trials that exploits the SL associated with *CCNE1* amplification. In this study, we identified and validated the genetic interaction between *FBXW7* and *PKMYT1,* proposing mutations in *FBXW7* as a novel predictive biomarker for lunresertib sensitivity. Importantly, several tumor types including colorectal and endometrial frequently carry nonsense truncating and hotspot missense (Arg465, Arg479, and Arg505) mutations in the *FBXW7* coding sequence at comparable proportions^46^. Hotspot mutations are mostly mono-allelic and translate to single amino acid changes in the WD40 repeats present in the C-terminus of FBW7, essential for substrate recognition. Both nonsense truncating and missense hotspot genetic alterations in *FBXW7* are believed to impair substrate ubiquitylation, and thus decreased degradation of oncogenes such as CCNE1, c-Myc, c-Jun, and Notch^46^. While traditional CRISPR-Cas9–based screens are well suited to study LoF truncating mutations, clinically relevant variants such as specific nonsense, missense, splicing, or silent mutations, exemplified by *FBXW7* hotspots, remain inaccessible to CRISPR-Cas9 high-throughput functional genomic approaches. Here, based on previous approaches^60^, we used base editing screens to identify mutations in *FBXW7* that sensitize cells to lunresertib. In addition to the known FBW7 Arg505 hotspot mutation, we identified five variants of unknown significance (His420, Arg441, Ser476, Asp487, and Gly644) as possible sensitizers to PKMYT1 inhibition. Importantly, once validated in preclinical models, these variants could guide patient enrollment in clinical trials of PKMYT1 inhibitors. A key limitation of this study was the restricted number of mutations accessible to cytosine base editors, mainly due to PAM sequence constraints and codon-change limitations. It is noteworthy that codons Arg465 and Arg479, frequently mutated in cancer, are not amenable to base editing under this setup and therefore are likely false negatives. Next-generation technologies, such as PAM-flexible base editors, adenine base editors, and prime editing, would overcome these constraints, enabling the testing of virtually any oncogenic variant^61^. Nevertheless, our data suggests that both truncating and mono-allelic hotspot mutations in *FBXW7* engender an equal vulnerability to *PKMYT1* genetic loss or pharmacological inhibition.

Here, we also showed the validation of a SL interaction between *CCNE1*-amplification and the two E2 ubiquitin conjugating enzymes of the APC/C, UBE2C and UBE2S. Since other subunits of the APC/C E3 ligase complex were also hits in our CRISPR screens, including *ANAPC7* and *ANAPC15*, we believe that this dependency could be generalized to the APC/C complex, and not an independent function of the E2 enzymes. APC/C activity drives timely cell cycle progression by relying on the co-activators CDC20 and CDH1^62,63^. While CDC20 is associated with the APC/C during early mitosis, CDH1 is associated with the APC/C during anaphase and interphase after cell division^64,65^. It was reported that Cyclin E1-CDK2 can phosphorylate CDH1, inhibiting the ability of APC/C^CDH1^ to degrade its substrates in G2 and early mitosis, including cyclin B1^66^. Interestingly, we previously found that *CCNE1*-overexpression activates the MMB–FOXM1 transcriptional program which also increases the abundance of mitotic substrates in S and G2 phase^13^. Thus, we propose that *CCNE1-UBE2C/S* SL interaction arises from the combined effect of elevated CCNE1-driven cell cycle progression and impaired mitotic timing control by the APC/C complex. It will be important for future work to determine the exact mechanism by which *CCNE1*-overexpressing cells are dependent on the activity of the APC/C E3 ubiquitin ligase.

Collectively, this work expands the map of SL interactions between tractable targets and prevalent oncogenic alterations in cancers with significant unmet clinical need. Moreover, our findings further support the use of CRISPR-based screens in isogenic cellular models for the identification of therapeutic targets. Finally, we highlight opportunities for targeted therapy development such as PKMYT1 (for *FBXW7*), UBE2C/S (for *CCNE1*-amplified), GFPT1 (for *ARID1A)*, and MARK2 kinase (for *STK11*) and provide a comprehensive resource for future studies to identify, validate, and characterize cancer driver–specific dependencies.

## Limitations of the Study

The study utilized CRISPR screens in cultured cell lines to identify SL interactions with common cancer driver genes. CRISPR screens have revolutionized the identification of SL interactions. However, several limitations of this technology need to be considered when interpreting the data. First, CRISPR screens can have false negative hits due to inefficient Cas9 editing activity which can significantly vary from cell line to cell line. Second, CRISPR screens can also have false negatives due to functionally redundant gene paralogs that can genetically compensate for each other. Importantly, genetic screens in cultured cell lines can yield false negative and positive hits due to two-dimensional growth on plastic using media with non-physiological growth factor composition and concentration. *In vivo* validation in animal models can mitigate the risk of SL interactions not translating to clinical settings.

## Supporting information

Table 1: Isogenic and DepMap SL screen data

Supplementary Table 1: Description of cell line engineering for each isogenic model.

Supplementary Table 2: Cancer cell lines used for DepMap synthetic lethal target identification.

Supplementary Table 3: PKMYT1i chemical genetic CRISPR screen in COL-hTERT Cas9 p53-/- cell line.

Supplementary Table 4: PKMYT1i cytosine base editing screen for FBXW7 in DLD-1 FNLS cell line.

Supplementary Table 5: List of all cell lines and culture conditions.

Supplementary Table 6: List of all sgRNAs and primers used for gene editing efficiencies.

Supplementary Table 7: Gene editing efficiencies for all CRISPR-based experiments.

Key Resource Table

## Resource Availability

### Lead Contact

Further information and requests for resources and reagents should be directed to and will be fulfilled by Jordan Young and Alejandro Álvarez-Quilón.

### Materials availability

Materials included in this manuscript will be shared upon request.

### Data and code availability

Raw sgRNA read counts for each CRISPR screen conducted in this study were deposited in Mendeley Data: https://doi.org/10.17632/k6wm46g4tw.1

## Acknowledgements

We would like to thank Ronny Drapkin for sharing the FT282-hTERT cell line; Jason Moffat for reagents and TKOv3 technical assistance; Traver Hart for guidance in the use of BAGEL2; Luke Dow and Raquel Cuella-Martin for technical advice on base editing.

## Author Contributions

Conceptualization, investigation, writing, and visualization, J.D., J.B.; formal analysis, data curation, and visualization, J.B.; investigation, C.B., S.J., T.G.R., D.G., M.E.O., N.L., A.R., J.M., J.L., V.B., L.L., A.L., J.H.L., M.C.M.; conceptualization, supervision, C.F., A.V., A.R., M.Z., D.D., S.J.M., M.Z.; conceptualization, supervision, writing, A.A.Q., J.T.F.Y.

## Supplemental Information

**Supplementary Figures 1-8**

**Supplementary Table 1:** Description of cell line engineering for each isogenic model.

**Supplementary Table 2:** Cancer cell lines used for DepMap synthetic lethal target identification.

**Supplementary Table 3:** PKMYT1i chemical genetic CRISPR screen in COL-hTERT Cas9 *p53^-/-^* cell line.

**Supplementary Table 4:** PKMYT1i cytosine base editing screen for *FBXW7* in DLD-1 FNLS cell line.

**Supplementary Table 5:** List of all cell lines and culture conditions.

**Supplementary Table 6:** List of all sgRNAs and primers used for gene editing efficiencies.

**Supplementary Table 7:** Gene editing efficiencies for all CRISPR-based experiments.

## STAR★METHODS

### Experimental Models

All the information about cell lines and animal models can be found in the Key Resource Table as well as in the Supplementary Tables 1 and 5.

## Method Details

### Patient population estimates for cancer drivers

The prevalence of each lesion was estimated using the TCGA PanCancer Atlas^67^ data sets on cBioPortal. The following Onco Query Language (OQL) queries were made in cBioPortal and downloaded as individual files for each genetic lesion: AMP (*CCNE1*), MUT=DRIVER or HOMDEL (*FBXW7, CDK12, ARID1A, KMT2D, TET2, KEAP1, STK11*), MUT=DRIVER (*DNMT3A, IDH1, SF3B1, SRSF2, U2AF1*) and HETLOSS (*SMAD4, RB1*). *SMAD4* was used as a surrogate for Chr18q loss and *RB1* was used as a surrogate for Chr13q loss. For each genetic lesion and cancer type, the percent of patients harboring the genetic lesion within that cancer type was multiplied by the estimated number of patients in the U.S. in 2024 that were diagnosed with that cancer type based on statistics from the American Cancer Society^68^.

### Cell culture

Cell lines were maintained at 37 °C and 5% CO_2_. Cell lines were authenticated by their respective vendors. The absence of mycoplasma contamination was regularly monitored using the MycoAlert Mycoplasma Detection kit (Lonza LT07-218). Culture conditions for all cell lines can be found in Supplementary Table 5. RPE1-hTERT Cas9 *p53^-/-^* cell line was described previously^69^. The PAC gene, encoding puromycin N-acetyltransferase, was knocked out by sgRNA nucleofection in RPE1-hTERT Cas9 *p53^-/-^* cells using the Lonza 4D-Nucleofector system according to the manufacturer’s instructions. Single PAC knockout clones were isolated by limiting dilution and validated by Sanger sequencing and puromycin sensitivity. 4D-Nucleofector programs for each cell line can be found in Supplementary Table 5. All sgRNA and primer sequences can be found in Supplementary Table 6. COL-hTERT Cas9 *p53^-/-^* PAC KO was generated by stable transduction of pLenti-Cas9-2A-Blasticidin and selection in media supplemented with 5 µg/mL blasticidin for 5 days. The PAC gene was then knocked out by sgRNA nucleofection followed by single clone isolation and validation by Sanger sequencing and puromycin sensitivity. To generate a *p53* knockout, *p53* sgRNA #4 was nucleofected and single clones were isolated by limiting dilution. Knockout of *p53* was confirmed by Sanger sequencing and p21 and p53 immunoblotting following treatment with Nutlin-3a at 5 µM for 36 h at 37 °C. Isogenic cell lines harboring knockout or knock-in mutations were generated by nucleofection of Cas9/sgRNA ribonucleoprotein complexes and single clones were isolated by limiting dilution. Cell lines were validated by Sanger sequencing and immunoblotting (Figure S2-4). sgRNA and ssODN sequences as well as PCR primers used for Sanger sequencing can be found in Supplementary Table 6. Antibodies used for immunoblotting are listed in the Key Resource Table. The A549 *KEAP1*-overexpressing cell line was generated by transduction of lentiviruses produced using pHIV-NAT-T2A-hCD52-KEAP1 and selected for 5 days with 1.6 mg/mL of nourseothricin. Again, single clones were isolated by limiting dilution and validated by immunoblotting. DLD-1 FNLS c9 was generated by clonal selection upon lentiviral infection with a FNLS-CBE3 expressing vector^70^.

### Antibodies, plasmids, and chemical compounds

See Key Resource Table.

### Lentivirus production and cell transductions

Lentiviral stocks were prepared by seeding 18 × 10^6^ Lenti-X 293T cells on a 15 cm dish in 20 mL media and transfected 24 h later by adding 2.4 mL of transfection mix, containing OptiMEM media, 18 μg of sgRNA plasmid, lentiviral packaging vectors (11.7 μg psPAX2 + 6.3 μg psPAX2), 90 μL Lipofectamine (Thermo Fisher Scientific) and 60 μL PLUS Reagent (Thermo Fisher Scientific). Medium was exchanged ∼16–20 h after transfection and supplemented with 20 mM HEPES. Virus-containing supernatant was collected ∼48 h post transfection, cleared by centrifugation at 1,000 x g RPM for 5 min and stored at −80 °C. Cells were transduced with the virus at a multiplicity of infection (MOI) of ∼0.5-1 in media supplemented with 1-5 µg/mL polybrene, depending on parental cell line sensitivity. Virus was removed 24 h post-infection and cells were selected with antibiotic selection media for the appropriate time.

### Gene editing analysis by genomic DNA extraction, PCR, and Sanger Sequencing

Cells were pelleted by centrifugation post-gene editing and genomic DNA was extracted using Qiagen DNeasy blood and tissue kits according to the manufacturer’s instructions. Primers were designed and synthesized to PCR-amplify genomic DNA surrounding the sgRNA site. All sgRNA and primer sequences can be found in Supplementary Table 6. PCR products were purified using Qiagen QIAquick PCR purification kit according to the manufacturer’s instructions. Purified PCR products were Sanger sequenced followed by gene editing sequence analysis using ICE software (https://ice.editco.bio/). Both genome-edited cell populations and single clones were analyzed using this method.

### CRISPR screens

Pooled lentiviral genome-wide CRISPR-KO screens were performed using protocols derived from literature^71^. Isogenic pairs were transduced with the lentiviral TKOv3 sgRNA library at a MOI of approximately 0.3. Puromycin-containing medium was added 2 days after infection. Four days after transduction (initial time point; t_0_), cells were pooled together and divided into two sets of technical replicates and grown for 18 days maintaining a library coverage of ≥400 cells per sgRNA at every step. Genomic DNA was isolated using QIAamp Blood Maxi Kit (Qiagen) and sgRNA sequences were amplified by PCR using NEBNext Ultra II Q5 Master Mix (New England Biolabs). The i5 and i7 multiplexing barcodes were added in a second round of PCR, and final gel-purified products were sequenced on an Illumina NextSeq500 system at the Lunenfeld-Tanenbaum Research Institute NBCC Facility (https://nbcc.lunenfeld.ca/) to determine the sgRNA representation in each sample. The screens were analyzed using BAGEL2^27^ and CCA^28^.

The RP-6175 chemical genetic CRISPR screen was performed as previously described^71^ but with several minor modifications. RP-6175 or DMSO treatment was added at t_6_ at a concentration of 200 nM and replenished every 4 days until completion of the screen at t_18_. Raw read counts from the t_18_ timepoint from the treatment condition were compared to DMSO control and analyzed with CCA^28^ and DrugZ^72^.

The base editing *FBXW7* CRISPR screen was performed following a previously described protocol^60^. Briefly, DLD-1 cells stably expressing the FNLS^47^ cytosine base editor were transduced with lentivirus carrying a library of sgRNAs tiling the *FBXW7* open reading frame (ENSG00000109670; CCDS3777: covering all exons and exon-intron junctions) at a MOI of ∼0.3. The screen was conducted in technical triplicates and a library coverage of >1,000 cells per sgRNA was maintained at every step. Puromycin-containing medium (4 µg/ml) was added 2 days after infection to select for transductants. Selection was continued until 96 h after infection, which was considered the initial time point (*t*_0_). RP-6306 was added to the cells at day 6 (*t*_6_) and 10 (*t_10_*) at 300 nM and the screen was terminated at *t*_18_. Sample processing was carried out as outlined above. FASTQ files were aligned to the library reference sequences using Bowtie^73^ to generate read counts for each sample replicate at the guide level. Raw read counts from the *t_18_* timepoint of each treatment condition were compared to DMSO control at *t_18_* using DESeq2^74^ to calculate log_2_ foldchange values, and the PCR2 forward primer used for each sample replicate was modeled as a covariate. The library of sgRNAs targeting FBXW7 was created using the crisprVerse^19^ package in R. The base editor object was assigned weights for each cytosine within the base editing window (positions -19 to -11 relative to the PAM) based on the FORECasT-BE model developed by Pallaseni et al. ^75^. Each sgRNA was then annotated for its most probabilistic predicted editing outcome at the amino acid level based on scores produced by crisprVerse^76^. Generally, all editing events with a score of at least 0.1 were considered, as well as the most likely event. Editing events that led to the same AA sequence were combined to produce scores and identify the most likely change. Splice changes were predicted by exact matches to the most common splice junction sequence at position -1 of the intron: AG on the forward strand or GT on the reverse strand.

### Synthetic lethal target identification using DepMap tumor cell line screens

Genomic data from the DepMap 2024Q2 public dataset^11^ was used to annotate cell lines and sort them into the ‘test’ group (cell lines positive for genetic lesion) or the ‘control’ group (cell lines wild-type for the genetic lesion). A summary of cell lines included in each test group and control group can be found in Supplementary Table 2, along with the criteria used for inclusion in ‘test’ and ‘control’ groups. The Wilcoxon rank-sum test was then used to compare DepMap Chronos^34^ scores across all genes between the two groups and generate nominal p-values, which were then adjusted using the Benjamini-Hochberg method^77^.

### Clonogenic survival assays and IncuCyte cell proliferation assays

Cells were seeded in 6-well plates, see Supplementary Table 5 for the appropriate seeding densities and assay lengths. Single cells were grown out until distinct colonies formed with greater than 50 cells per colony. Colonies were rinsed with PBS and stained with 0.4% (w/v) crystal violet in 20% (v/v) methanol for 30 min. The stain was aspirated, and plates were rinsed twice in double-distilled H2O and air-dried. Colonies were counted using a GelCount instrument (Oxford Optronix, GelCount).

For IncuCyte proliferation assays, cells were seeded in 96-well plates, also see Supplementary Table 5 for the appropriate seeding densities and assay lengths. Cell confluency was monitored once or twice a day until the untreated cells were confluent using an IncuCyte S3 Live-Cell Imager (Sartorius). Drugs were added 24 h after seeding using Tecan D300E dispenser and refreshed every 3–4 days.

### Immunoblotting

Cell pellets were extracted by incubation in RIPA lysis and extraction buffer (Thermo Scientific, 89900) with added Halt Protease inhibitor cocktail (Thermo Scientific, #78410) and Halt Phosphatase inhibitor cocktail (Thermo Scientific, #78420) on ice for 20 min followed by centrifugation at 14000 x g for 15 min at 4 °C. Cleared cell lysates were then quantified using BCA assay (Thermo Scientific, 23225). Protein extracts were diluted in 2x sample buffer (Thermo Scientific, LC2676) and boiled for 10 min prior to separation by SDS-PAGE on Novex Tris-Glycine gradient gels. Proteins were transferred onto nitrocellulose membranes (ThermoFisher, PB7220) in 1x Novex Tris-Glycine Transfer Buffer (ThermoFisher, LC3675) containing 20% methanol and 0.04% SDS. Membranes were blocked in 5% milk/Tris-buffered saline/0.1% Tween 20 (5% milk/TBST) and incubated with primary antibodies diluted 1:1000 in 5% milk/TBST either overnight at 4°C or for 2 h at room temperature (RT). Membranes were then washed 3x 5min with TBST and incubated with a secondary antibody 1:5000 for 1 h at RT, after which they were washed again and developed using the Pierce ECL Pico or Femto Western Blotting substrate (Thermo Fisher Scientific). Chemiluminescence was detected using iBright FL1500 imaging system (Thermo Fisher).

### RNA extraction, cDNA synthesis, and quantitative PCR

RNA was extracted from harvested cell pellets using RNeasy Plus Mini Kit (Qiagen) according to the manufacturer’s protocol. RNA was eluted in RNAse/DNAse free water and measured with Nanodrop (Thermo). cDNA was synthesized from 1 µg of RNA using High-Capacity cDNA reverse transcription kit (Thermo) and random primers. Quantitative PCR was performed using TaqMan™ Fast Advanced Master Mix (Applied Biosystems) in a QuantStudio 7 Flex (Applied Biosystems) with TaqMan Assay probes (ARID1A: Hs00195664_m, GFPT1: Hs00899865_m1, GFPT2: Hs01049570_m1, PPIA: Hs04194521_s1). Relative expression to PPIA was reported and normalized on BEAS-2B WT.

### Phospho-H3 (Ser10) flow cytometry assay

Cells were pulsed with 20 μM EdU (Life Technologies cat. no. A10044) for 30 min, collected by trypsinization, resuspended as single cells, washed once in PBS and pelleted at 600g for 3 min at 4 °C. All subsequent centrifugations were performed in this manner. Cells were fixed in 4% PFA/PBS for 15 min at RT and excess ice cold PBSB (1% BSA in PBS, 0.2 μM filtered) was added before pelleting. Cells were resuspended in permeabilization buffer (PBSB, 0.5% Triton-X 100) and incubated at room temperature for 15 min. Excess blocking buffer (PBSB, 0.1% NP-40) was added, cells were pelleted, resuspended in blocking buffer containing primary antibodies and incubated at room temperature for 1 h. Excess blocking buffer containing secondary antibodies was added, cells were pelleted, resuspended in blocking buffer and incubated at room temperature for 30 min. Excess blocking buffer was added, cells were pelleted and washed one additional time in PBSB. Cells were resuspended in EdU staining buffer (150 mM Tris-Cl pH 8.8, 1 mM CuSO4, 100 mM ascorbic acid and 10 μM AlexaFluor 555 azide (Life Technologies, cat. no. A20012)) and incubated at room temperature for 30 min. Excess PBSB was added, cells were pelleted and washed one additional time in PBSB. Cells were resuspended in analysis buffer (PBSB, 0.5 µg ml−1 DAPI, 250 µg µl−1 RNase A (Sigma-Aldrich, cat. no. R4875)) and incubated at 37 °C for 30 min or left at 4 °C overnight. Cells were analyzed at the LTRI flow cytometry facility on a Fortessa X-20 (Becton Dickinson) using FACSDIVA v8.0.1 with at least 9,000 events collected and analyzed using FlowJo v10.

### Nuclear phospho-cyclin B (Ser126) and γH2AX QIBC assays

3,000 cells per well were seeded in 96-well plates and cultured for up to 72 h depending on the experiment. Before collection, cells were pulsed with 20 μM EdU (5-ethynyl-2-deoxyuridine, Life Technologies cat. no. A10044) for 30 min and then washed with PBS and fixed with 4% paraformaldehyde (PFA) in PBS for 15 min at room temperature. Cells were then rinsed with PBS and permeabilized using 0.3% Triton X-100/PBS for 30 min. Cells were washed with PBS and incubated in blocking buffer (10% goat serum (Sigma cat. no. G6767), 0.5% NP-40 (Sigma-Aldrich, cat. no. I3021), 5% w/v saponin (Sigma-Aldrich, cat. no. 84510), diluted in PBS) for 30 min. Fresh blocking buffer containing primary antibodies was added for 2 h. Cells were rinsed three times with PBS and then blocking buffer, with secondary antibodies and 0.4 μg ml−1 DAPI (4,6-diamidino-2-phenylindole, Sigma-Aldrich, cat. no. D9542) was added for 1 h. After rinsing with PBS, immunocomplexes were fixed again using 4% PFA/PBS for 5 min. Cells were rinsed with PBS and incubated with EdU staining buffer (150 mM Tris-Cl pH 8.8, 1 mM CuSO4, 100 mM ascorbic acid and 10 μM AlexaFluor 555 azide (Life Technologies, cat. no. A20012) for 30 min. Wells were filled with 200 μl PBS and images were acquired at the Network Biology Collaborative Centre (LTRI) on an InCell Analyzer 6000 automated microscope (GE Life Sciences) with a 20× objective. Image analysis was performed using Cellprofiler 3.1.9 and RStudio v1.2.501957.

### Doxycycline-inducible shRNA mediated gene knockdown

Partial depletion of GFPT1 was performed by the introduction of two doxycycline-inducible GFPT1 shRNAs by lentiviral transduction into the BEAS-2B *ARID1A* isogenic pair. Briefly, 200,000 cells were seeded in a 6-well plate in polybrene-supplemented media (5 µg/mL) and transduced with SMARTvector inducible lentiviral shRNA particles (Horizon Discovery) at a MOI of about 1. The day following transduction, puromycin-supplemented media (2 µg/mL) was added to the cells for 48 h until selection was completed. Inducible shRNA-containing cell populations were then seeded for clonogenic survival assays and treated with 0.1, 0.5, or 1 µg/mL of doxycycline. Doxycycline-containing media was replenished every 3-4 days until colonies reached ∼6 population doublings. Similarly, each tumor cell line from assembled *ARID1A* wild-type and mutant panel was also transduced with SMARTvector inducible shRNA #183 lentiviral particles and seeded for clonogenic survival assays with 0, 0.1, or 0.2 µg/mL doxycycline.

### Cell line- and patient-derived xenografts

Animals were housed and experiments performed at Repare Therapeutics (adMare site, Saint Laurent, Quebec), which is a Canadian Council on Animal Care accredited vivarium. Studies were conducted under a protocol approved by the adMare Institutional Animal Care Committee (aIACC). Mice were inspected upon arrival and group-housed (3-5 per cage) in individual HEPA ventilated autoclaved cages (Innocage^®^ IVC, Innovive, San Diego, CA, USA) in a temperature-controlled environment (22±1.5 °C, 30-80 % relative humidity, 12 h light/dark). Animals were provided with autoclaved corncob bedding, irradiated food (Harlan Teklad, Montreal, Canada) and autoclaved water *ad libitum*. They were also provided with nesting material (Ketchum, cat#087) and a plastic shelter as enrichment. Fresh bedding, nesting material and water were replaced on a weekly basis. Mice were acclimatized in the animal facility for at least 5 days prior to use and were identified with indelible ink. Experiments were performed during the light phase of the cycle.

TOV-21G shRNA #183 GFPT1 (Clone4a) cells were implanted at 2×10^6^ cells per mouse into the right flanks of female NCG mice (5-7 weeks old; Charles River), in 1:1 matrigel: media (Matrigel Corning, cat# CB35248). When tumors reached the target size of 100-150 mm^3^, (*n* = 6) mice were randomized to treatment groups. After randomization, doxycycline was administered *i.p.* at 20 mg/kg once daily for the first 2 days and was also provided in chow at 2g/kg (Bio-serv, Cat #S10204) *ad libitum* for the duration of the experiment. Vehicle control mice received PBS *i.p.* and blank chow (Bio-Serv, Cat#S10203). Tumors were harvested from a cohort of satellite mice after 3, 6, and 9 days of doxycycline treatment and snap frozen in ceramic bead containing tubes (OMNI Cat# 19-628). Frozen tumor samples were then lysed in MSD Tris lysis buffer (MSD, Cat# R60TX-2) containing Halt Protease inhibitor cocktail (Thermo Scientific, #78410) and Halt Phosphatase inhibitor cocktail (Thermo Scientific, #78420) for downstream immunoblotting analysis.

In vivo studies using patient-derived xenografts (PDX) were conducted at Crown Biosciences Inc. Fresh primary human tumor tissue was harvested and cut into small pieces (approximately 2-3 mm in diameter). These tumor fragments were inoculated subcutaneously into the right flank of female BALB/c nude mice (5-7 weeks old) for tumor development and subsequently passaged by implantation into the cohort of mice enrolled in the efficacy study. Mice were randomized according to growth rate into treatment groups (*n* = 8) when the mean tumor size reached approximately 150 (100-200) mm^3^. The procedures involving the care and use of animals for the PDX models were reviewed and approved by the CrownBio Institutional Animal Care and Use Committee (IACUC) prior to execution. During the study, the care and use of animals were conducted in accordance with the regulations of the Association for Assessment and Accreditation of Laboratory Animal Care (AAALAC). RP-6306 was formulated in 0.5% methylcellulose and orally administered twice daily (*b.i.d.*, 0-8h) for up to 29 days. Animals were monitored for tumor volume, clinical signs and body weight twice weekly.

For all experiments, tumors were measured using vernier calipers and volumes calculated using the formula 0.52×L×W^2^. Response to treatment was evaluated for tumor growth inhibition (%TGI) defined as: % TGI= ((TV_vehicle/last_ – TV_vehicle/day0_) - (TV_treated/last_ – TV_treated/day0_)) / (TV_vehicle/last_ – TV_vehicle/day0_) x100 calculated based on the mean tumor volumes at day 0 and on the last day of measurement. According to aIACC and IACUC approved animal protocols, mice were sacrificed as soon as their tumor volume exceeded 2000 mm^3^. Change in body weight (BW) was calculated based on individual body weight changes relative to day 0 using the formula: % BW change = (BW_last_-BW_day0_/ BW_day0_) x100.

### D-2-Hydroxyglutarate (2-D-HG) quantification assay

Quantification of 2-D-HG was performed following a previously described protocol^72^. Briefly, L- α-hydroxyglutaric acid disodium salt and D- α-hydroxyglutaric acid disodium salt (Sigma) was used as calibration standards after derivatization with DATAN. Pellets from 10^6^ RPE1-hTERT Cas9 *p53^-/-^* cells were combined with an ice-cold solution of 80% methanol in water, vortexed and centrifuged. 20 μL of supernatant were collected and dried under a nitrogen stream. To the dried samples, 100 μL of 50 mg/mL DATAN, (+)-O,O’-diacetyl-L-tartaric anhydride in dichloromethane (Sigma) and mixed for 1 h at 75 °C. Finally, the samples were dried again and reconstituted with 1:1 acetonitrile:water containing an internal standard for LC-MS analysis. Separations were performed using an Acquity-H class UPLC equipped with an HSS T3 (2.1 x 100 mm, 1.8 μM) column from Waters. A gradient elution was employed using mobile phase A: 125 mg/L ammonium formate in water adjusted to pH 3.5 with formic acid against mobile phase B:methanol. Mass spectrometry detection was achieved using a Waters Xevo Qtof G2 operated in positive mode with an electrospray source.

## Supplementary Figure Legends

**Figure S1.**
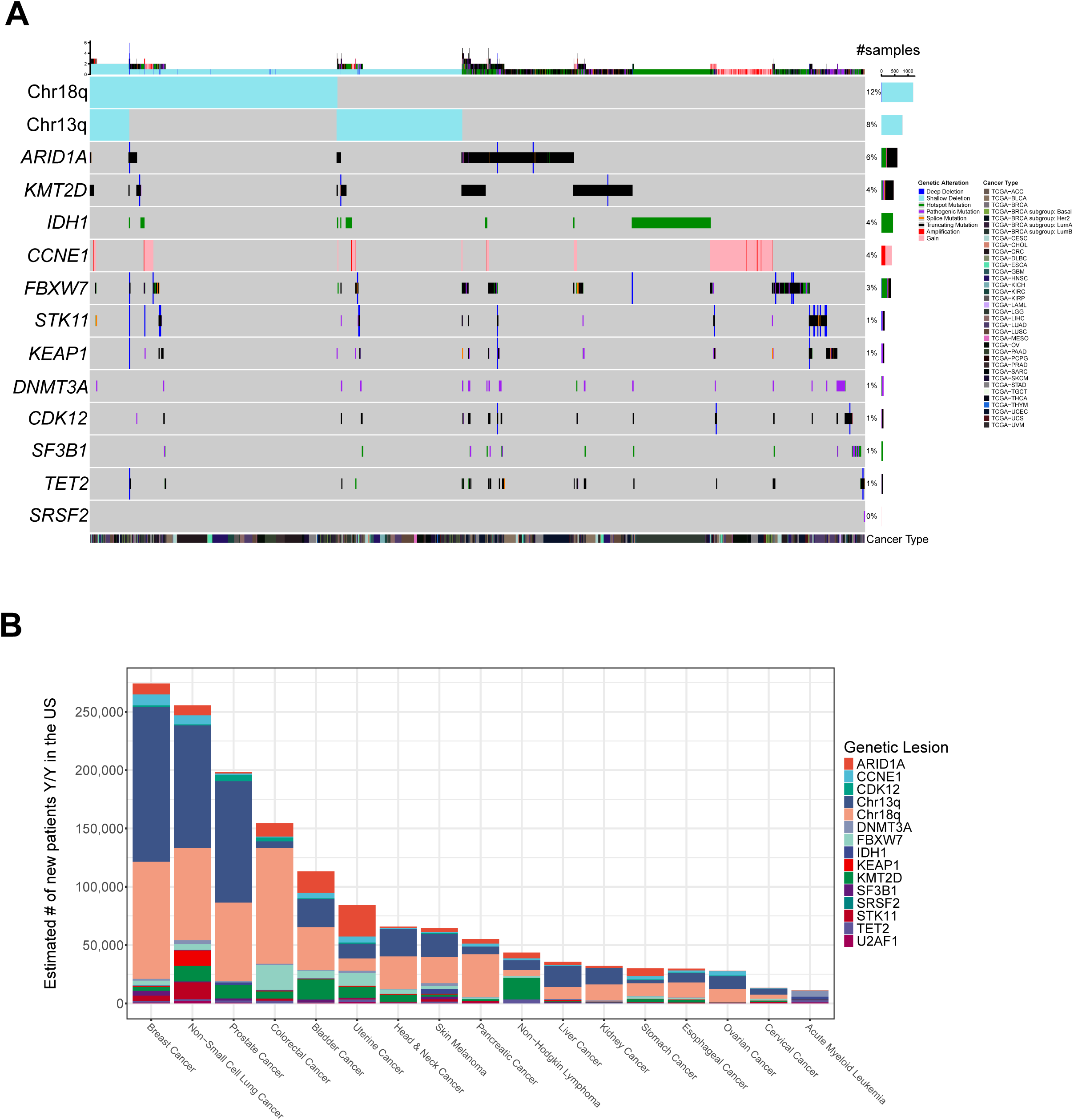
Landscape of prevalent cancer driver alterations analyzed in this study. **A)** OncoPrint of genomic driver alterations ranked by abundance of samples in the indicated TCGA cancer types. **B)** Representation of the estimated number of new patients per year with tumors harboring the indicated genetic alterations. Calculated using TCGA prevalence data and number of new diagnoses in 2024 from the American Cancer Society.

**Figure S2.**
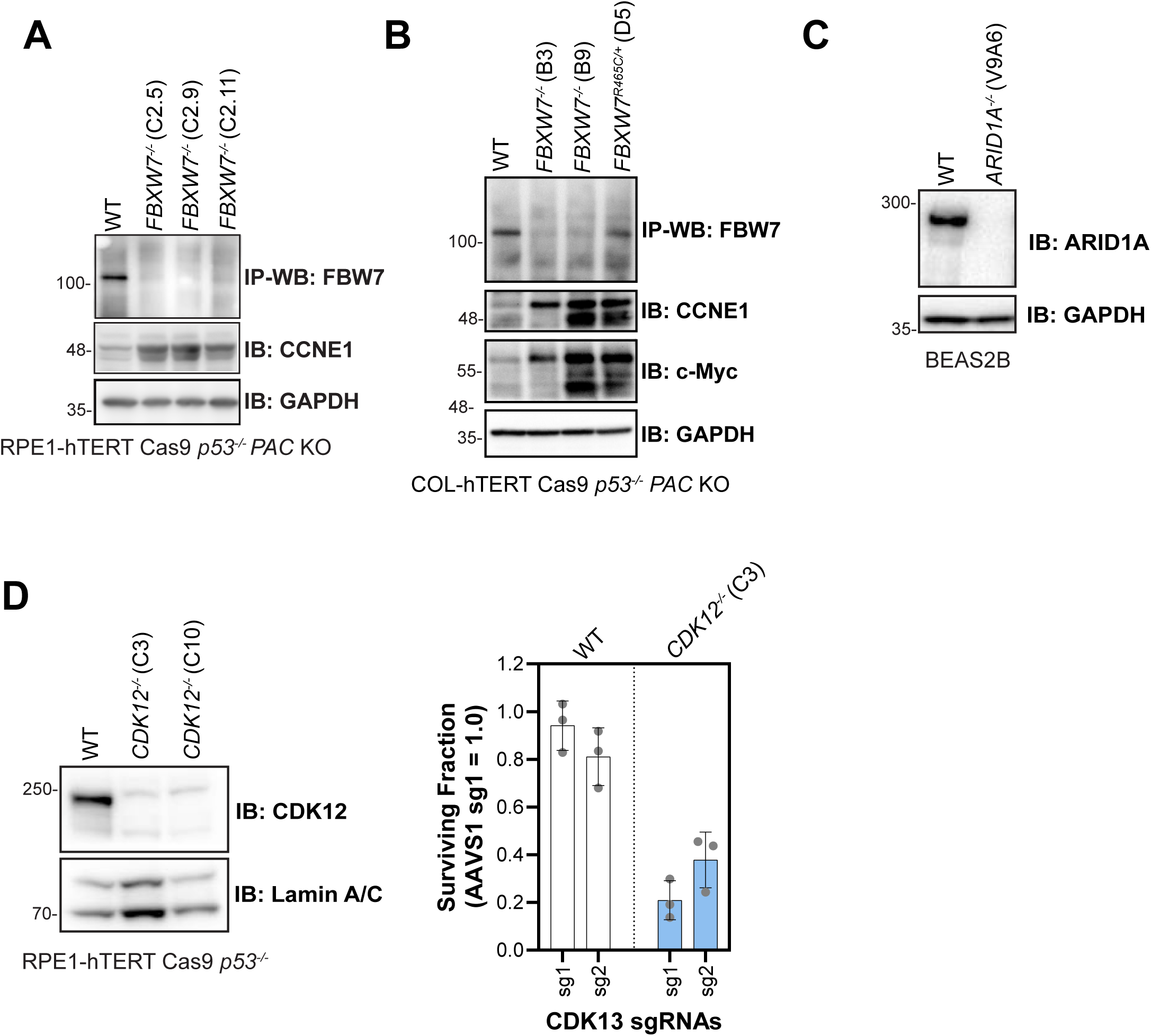
Characterization of *FBXW7*, *ARID1A*, and *CDK12* isogenic cell line models. **A)** Whole cell lysates or FBW7-immunoprecipitated samples of RPE1-hTERT Cas9 *p53^-/-^*with the indicated *FBXW7* genotypes were immunoblotted with FBW7, cyclin E (CCNE1), and GAPDH specific antibodies. **B)** Whole cell lysates or FBW7-immunoprecipitated samples of COL-hTERT Cas9 *p53^-/-^*with the indicated *FBXW7* genotypes were immunoblotted with FBW7, cyclin E (CCNE1), c-Myc, and GAPDH specific antibodies. **C)** Immunoblot using whole cell lysates of BEAS-2B wild-type or an *ARID1A* knockout clone (V9A6) with ARID1A and GAPDH specific antibodies. **D)** Left, immunoblot using whole cell lysates of RPE1-hTERT Cas9 *p53^-/-^* and two independent *CDK12* knockout clones with CDK12 and Lamin A/C specific antibodies. Right, clonogenic survival of RPE1-hTERT Cas9 *p53^-/-^* and *CDK12^-/-^* clone (C3) after introduction of sgRNAs targeting either *AAVS1* or the *CDK13* gene. Data are represented as the mean ± SD (*n* = 3).

**Figure S3.**
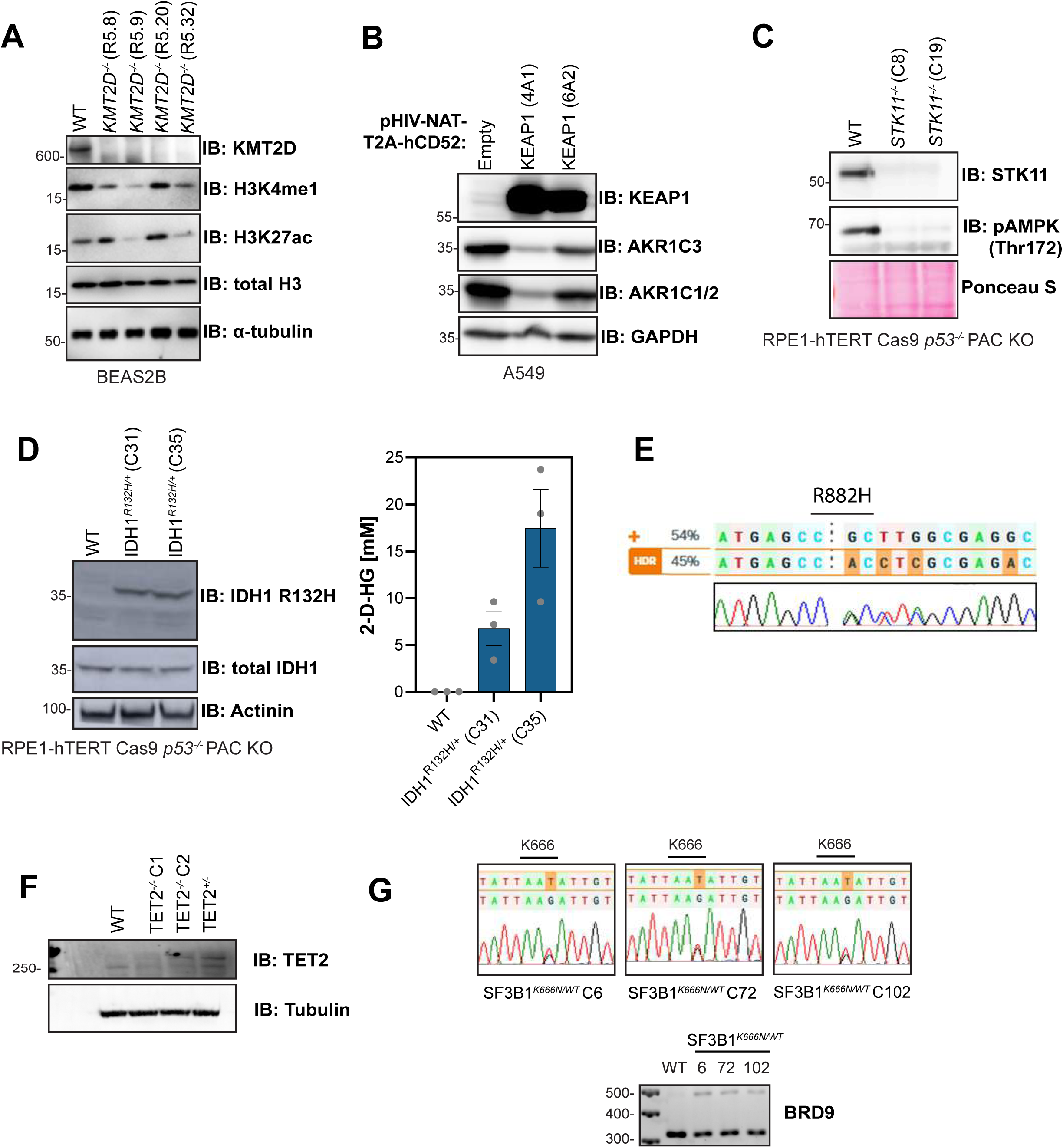
Characterization of *KMT2D*, *KEAP1*, *STK11*, *IDH1*, *TET2*, and *SF3B1* isogenic cell line models. **A)** Whole cell lysates of BEAS-2B wild-type and four independent *KMT2D* knockout clones were immunoblotted with KMT2D, H3K4me1, H3K27ac, histone 3 (H3), and α-tubulin specific antibodies. **B)** Immunoblots using antibodies against KEAP1, AKR1C1/2/3, and GAPDH from whole cell lysates of *KEAP1*-mutant A549 lung cancer cells and two independent clones with stable overexpression of wild-type *KEAP1*. **C)** STK11 and pAMPK (Thr172) immunoblots using whole cell lysates from RPE1-hTERT Cas9 *p53^-/-^* parental cells and two independent *STK11* knockout clones. **D)** Left, immunoblot using whole cell lysates of RPE1-hTERT Cas9 *p53^-/-^* parental cells and two independent *IDH1^R132H/+^*mutant clones using IDH1, mutant IDH1 (IDH1-R132H), and Actinin specific antibodies. Right, quantification of D-2-hydroxyglutarate (D-2-HG) oncometabolite in RPE1-hTERT Cas9 *p53^-/-^* cells with the indicated *IDH1* genotypes. **E)** Sanger sequencing chromatogram showing the outcome of *DNMT3A^R882H/+^* CRISPR knock-in in K562 cells. **F)** Whole cell lysates of RPE1-hTERT Cas9 *p53^-/-^* parental cells, two independent *TET2* knockout clones, and one heterozygous *TET2* knockout clone were immunoblotted with TET2 and α-tubulin specific antibodies. **G)** Top, Sanger sequencing chromatograms showing the outcome of *SF3B1^K666N/+^* CRISPR knock-in in three independent K562 heterozygous clones. Bottom, Agarose gel showing RT-PCR products from *BRD9*-specific primers in three K562 *SF3B1^K666N/+^*clones.

**Figure S4.**
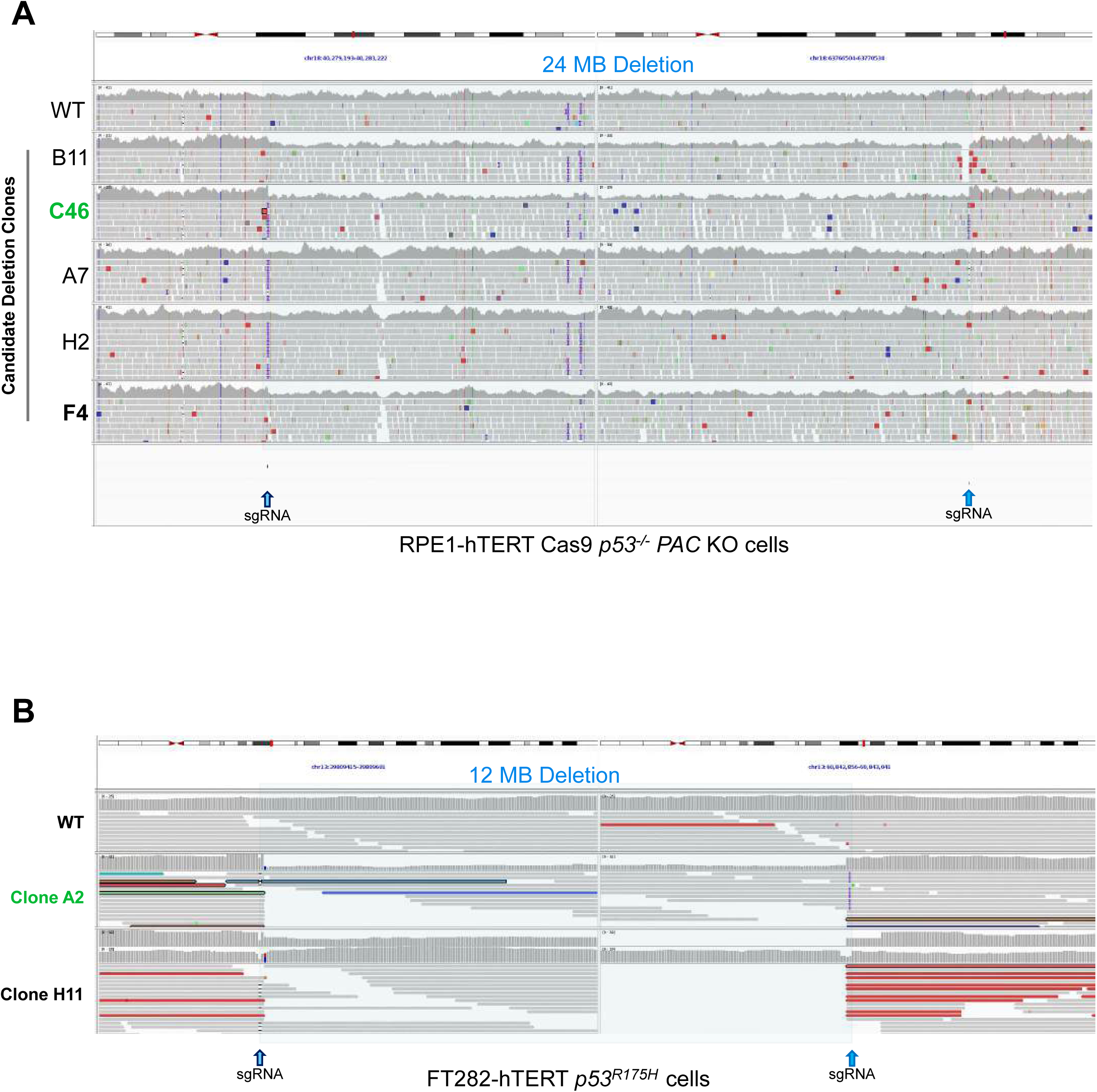
Characterization of hemizygous arm-level aneuploidy in Chr18q and Chr13q isogenic cell line models. **A)** Linear genome view of Chr18q showing whole genome sequencing alignments of RPE1-hTERT Cas9 *p53^-/-^*parental cells and five independent clones after simultaneous introduction of two sgRNAs (blue arrows). A hemizygous deletion of approximately 24 MB can be observed in clone C46. **B)** Linear genome view of Chr13q showing whole genome sequencing alignments of FT282-hTERT *p53^R175H^* parental cells and two independent clones after simultaneous introduction of two sgRNAs (blue arrows). A 12 MB hemizygous deletion can be observed in clone A2.

**Figure S5.**
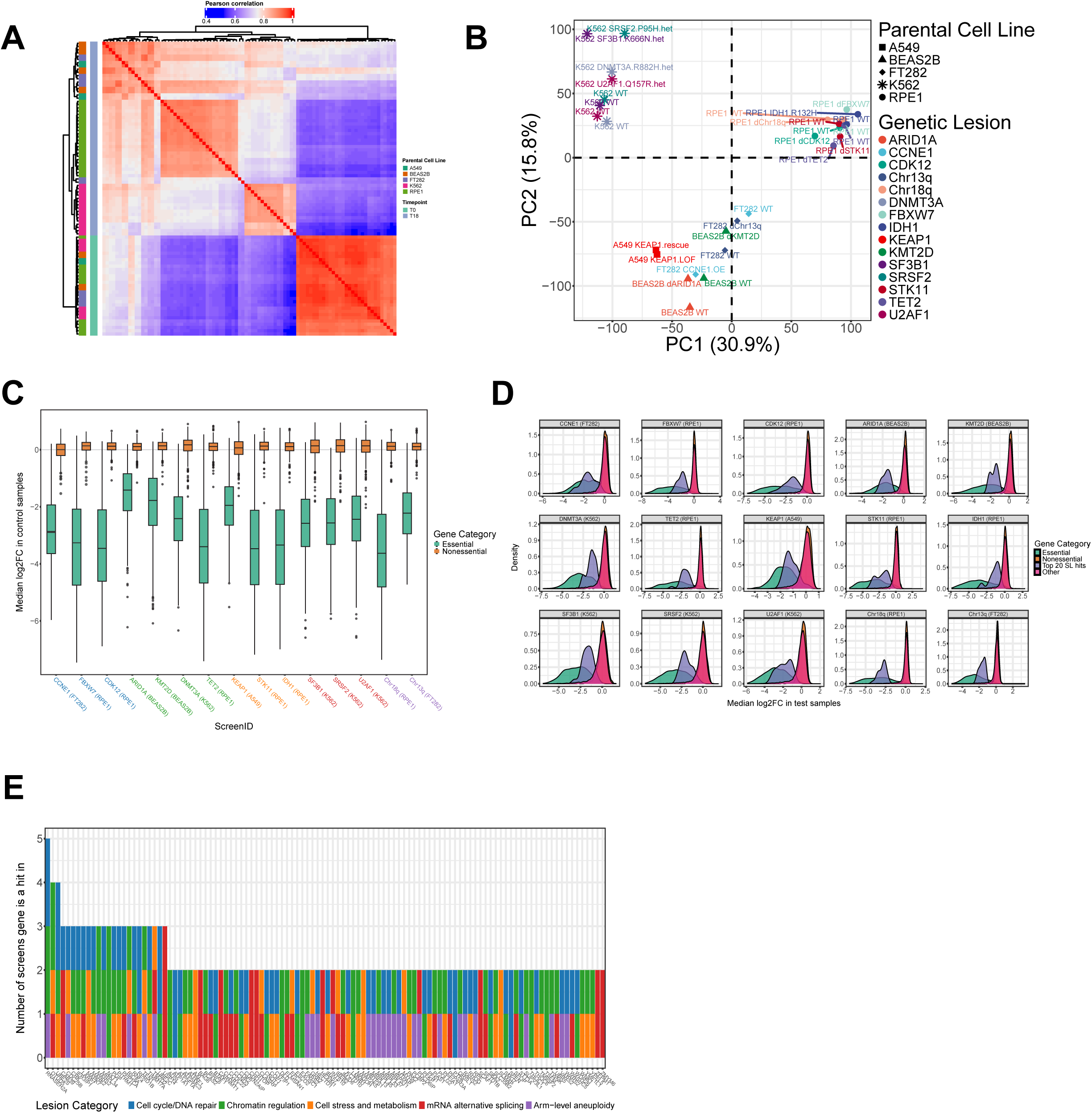
CRISPR screen quality control metrics. **A)** Heatmap showing clustering of samples by Pearson correlation of normalized sgRNA read counts. **B)** Principal component analysis (PCA) conducted using the median log2 fold-change values at the gene level. **C)** Boxplots showing dropout of essential gene sgRNAs vs. non-essential gene sgRNAs across all screens. **D)** Density plots showing the distributions of median log2 fold-change values for the top 20 SL hits, all essential genes, all non-essential genes, and the remainder of all genes across screens. **E)** Boxplot showing the number of times a given gene appeared in the top 50 hits across all screens. Boxes are colored by lesion category, and only genes appearing in the top hits of at least 2 screens are shown.

**Figure S6.**
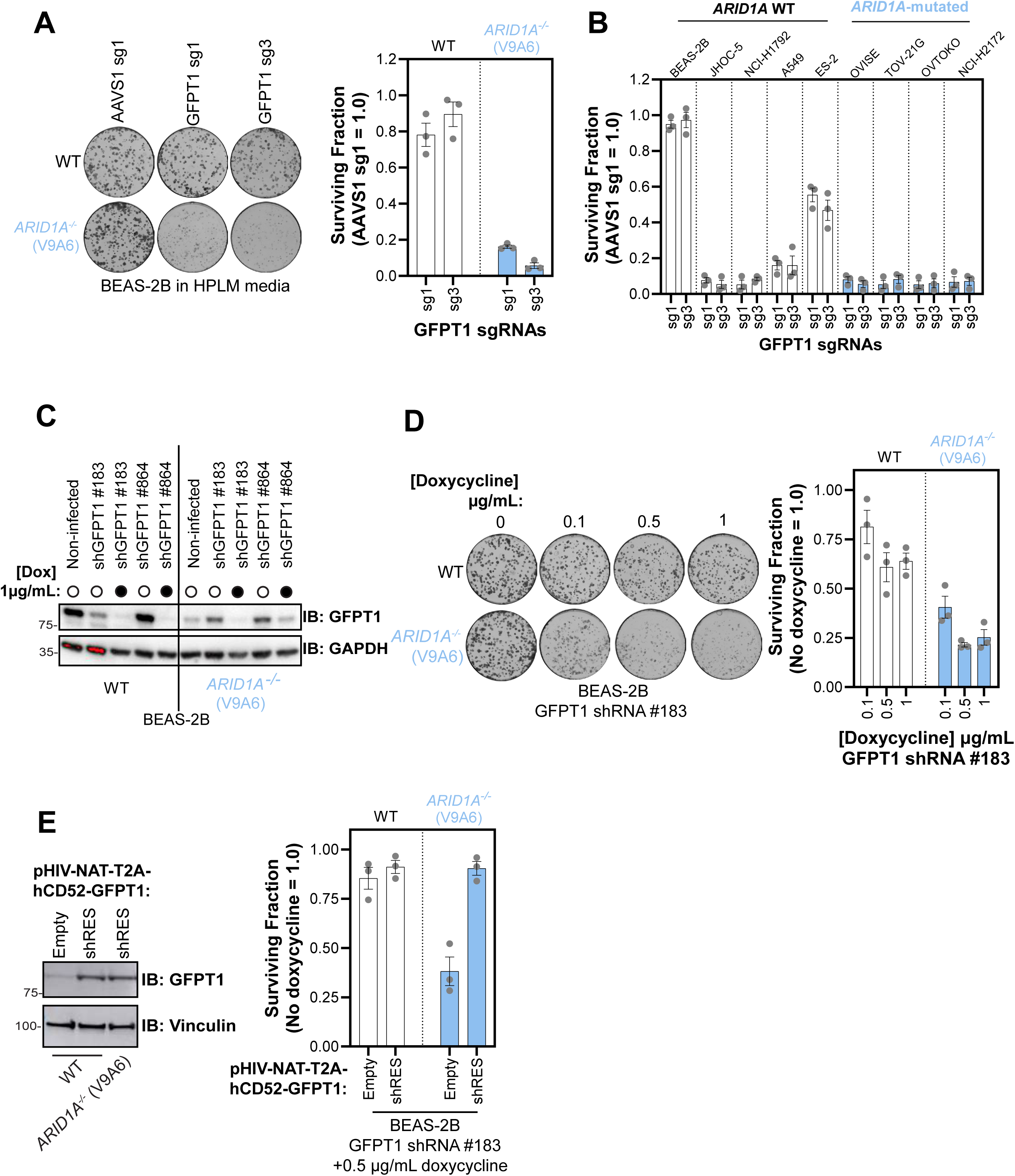
*GFPT1* is synthetic lethal with *ARID1A*. **A)** Clonogenic survival assays using human plasma-like media (HPLM) and BEAS-2B wild-type or *ARID1A^-/-^* cell lines after introduction of *AAVS1* or *GFPT1* sgRNAs. Left, representative images of colonies stained with crystal violet. Right, quantification of clonogenic survival assays. Data are represented as the mean ± SD (*n* = 3). **B**) Quantification of clonogenic survival assays in tumor-derived cell lines grouped according to their *ARID1A* status (wild-type or LoF). Cells were nucleofected with indicated *GFPT1* targeting sgRNA or the *AAVS1* targeting control sgRNA. Data are the mean ± SD (*n* = 3). **C**) Whole cell lysates of BEAS-2B wild-type and *ARID1A^-/-^* (clone V9A6) transduced with lentivirus expressing GFPT1 targeting shRNAs. After exposure to the indicated concentration of doxycycline or vehicle, lysates were immunoblotted with GFPT1 and GAPDH specific antibodies. **D**) Clonogenic survival assays of BEAS-2B wild-type and *ARID1A^-/-^* cells after doxycycline-induced expression of GFPT1 shRNAs. Left, representative images of colonies stained with crystal violet. Right, quantification of clonogenic survival assays. Data are represented as the mean ± SD (*n* = 3). **E**) Left, immunoblot using GFPT1 and vinculin specific antibodies of whole cell lysates from BEAS-2B wild-type and *ARID1A^-/-^* cells transduced with empty or a shRNA-resistant GFPT1 mutant open reading frame. Right, clonogenic survival assays in BEAS-2B wild-type and *ARID1A^-/-^* cells transduced with a doxycycline-inducible shRNA targeting endogenous *GFPT1* mRNA (shGFPT1#183) and also tranduced with lentiviral particles expressing a shGFPT1#183-resistant mutant GFPT1 open reading frame. Cells were exposed to 0.5 µg/mL of doxycycline to induce knockdown of endogenous *GFPT1*.

**Figure S7.**
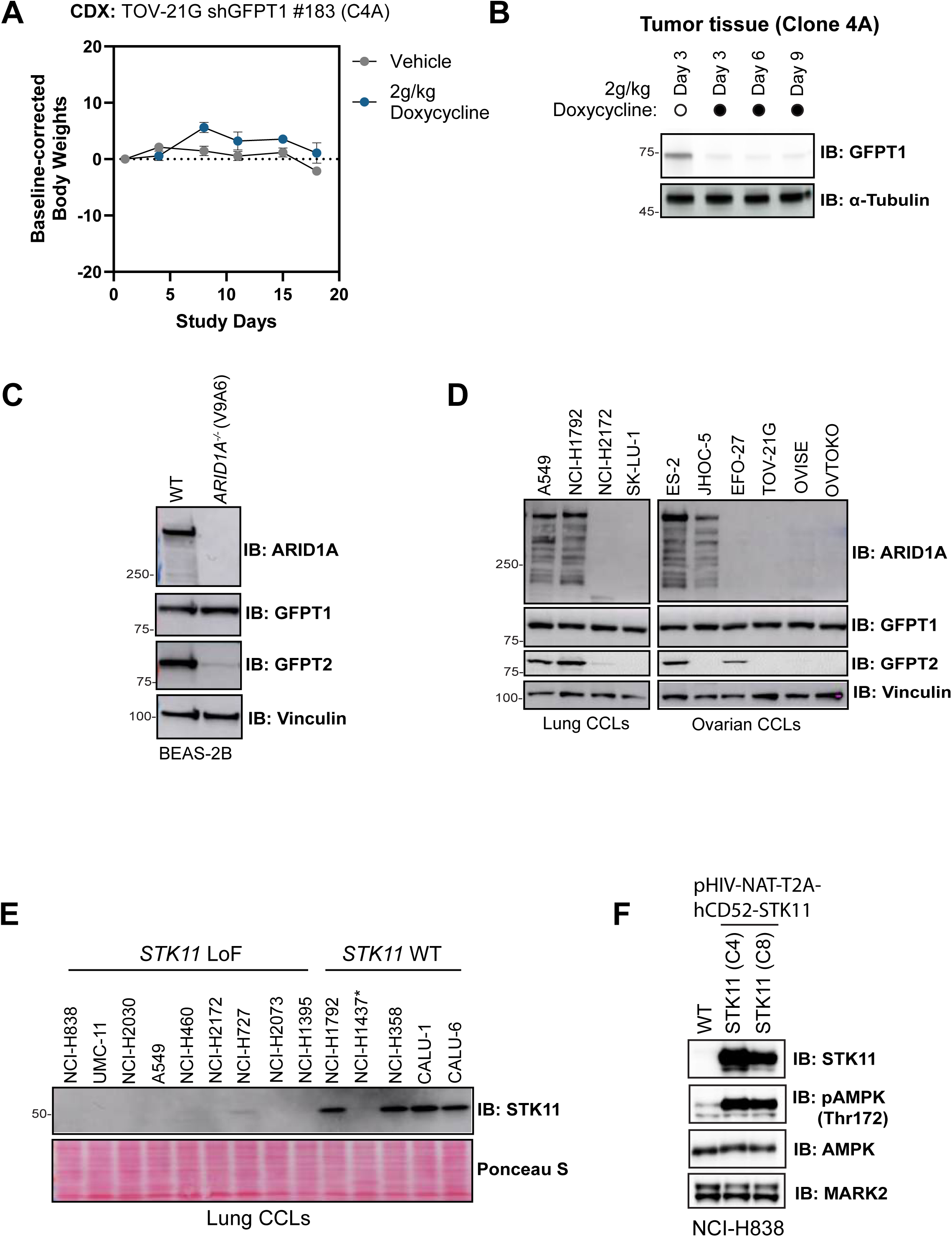
Additional validation of *GFPT1-ARID1A* and *MARK2-STK11* synthetic lethality. **A)** Body weight changes of mice bearing TOV-21G clone 4A (shGPT1 #183) cell derived xenografts and consuming chow containing 2g/kg of doxycycline. **B)** GFPT1 and α-tubulin immunoblots of TOV-21G CDX tumor lysates prepared at either 3, 6, or 9 days after mouse treatment with 2 g/kg of doxycycline or vehicle. **C)** Whole cell lysates of BEAS-2B wild-type or *ARID1A^-/-^* (clone V9A6) were immunoblotted with ARID1A, GFPT1, GFPT2, and vinculin specific antibodies. **D)** Whole cell lysates of the indicated lung and ovarian cancer-derived cell lines (CCLs) were immunoblotted with ARID1A, GFPT1, GFPT2, and vinculin specific antibodies. **E)** Whole cell lysates of the indicated lung cancer-derived cell lines (CCLs) were immunoblotted with a STK11 specific antibody. Ponceau S staining is shown as a loading control. **F)** Whole cell lysates from NCI-H838 wild-type parental and two *STK11-*overexpression clones were immunoblotted with a STK11, pAMPK (Thr172), AMPK, and MARK2 specific antibodies.

**Figure S8.**
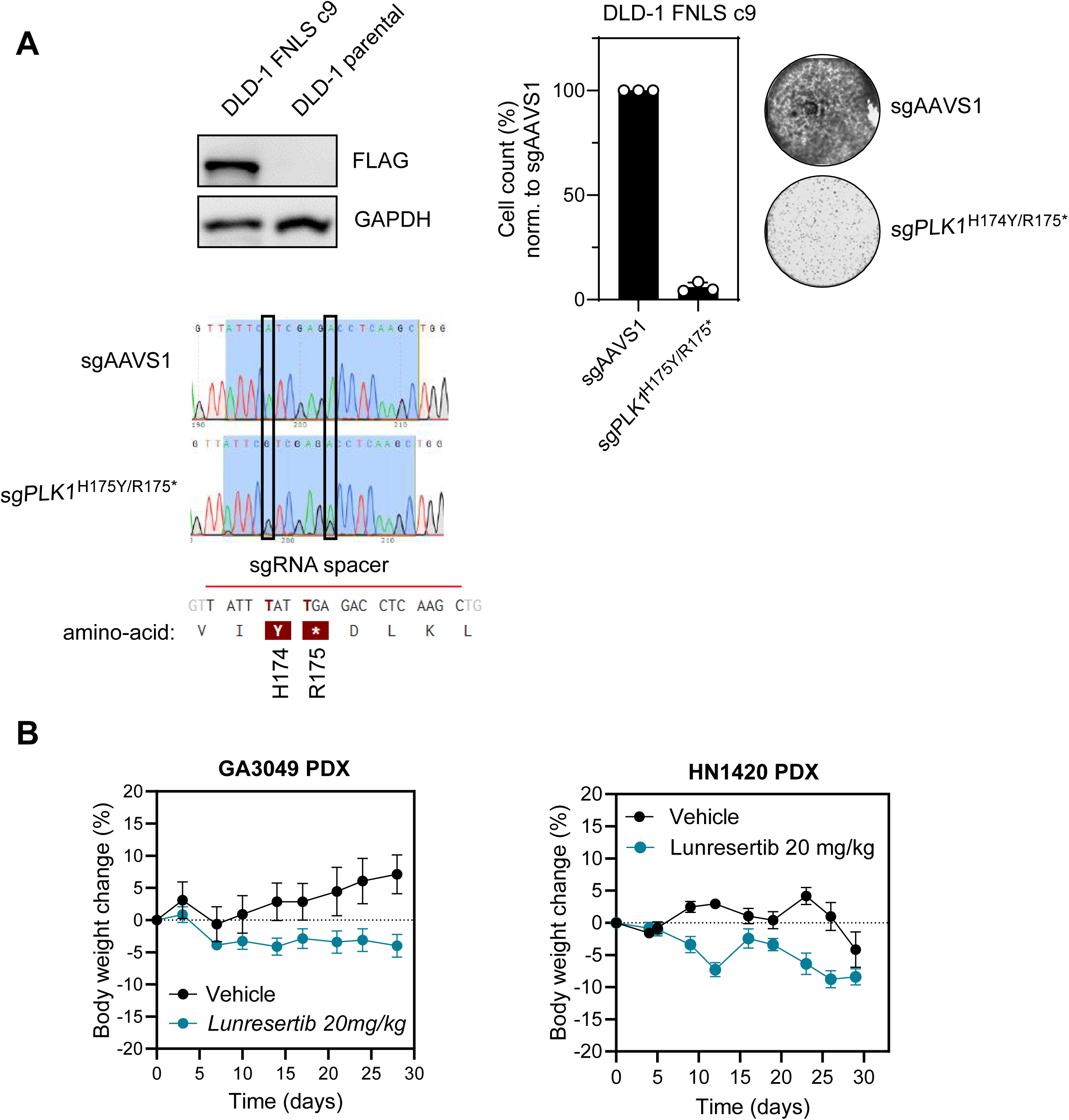
Characterization of DLD-1 base editing cell line and body weight changes for mice in the *FBXW7* PDX study. **A)** Left, whole cell lysates of DLD-1 parental and clone 9 transduced with a construct stably expressing a FLAG-tagged version of the FNLS cytosine base editor were immunoblotted with FLAG and GAPDH specific antibodies. Right, quantification of clonogenic survival and representative images of cells stained with crystal violet for DLD-1 parental and clone 9 electroporated with sgRNAs targeting AAVS1 or *PLK1* (introducing a nonsense truncating mutation by base editing). Bottom, Sanger sequencing chromatogram showing the outcome of cytosine base editing upon electroporation of *PLK1* targeting sgRNA or AAVS1 control. **B)** Body weight changes of mice bearing patient-derived gastric (left) and head and neck (right) tumors treated with 20 g/kg of lunresertib (RP-6306) or vehicle over time.

